# Identification of potent and orally efficacious phosphodiesterase inhibitors in s*Cryptosporidium parvum-*infected immunocompromised mice

**DOI:** 10.1101/2023.09.26.559556

**Authors:** Jose E. Teixeira, Makafui Gasonoo, Peter Miller, Jubilee Ajiboye, Alexandra C. Cameron, Erin Stebbins, Scott D. Campbell, David W. Griggs, Thomas Spangenberg, Marvin J. Meyers, Christopher D. Huston

## Abstract

*Cryptosporidium* species, mostly *C. parvum* and *C. hominis* in humans, are intestinal apicomplexan parasites that cause life-threatening diarrhea in young children and people with cell-mediated immune defects, such as due to AIDS. There is only one approved treatment for cryptosporidiosis, but it is ineffective for immunocompromised people and only modestly effective for children. In this study, screening 278 compounds from the Merck KGaA, Darmstadt, Germany collection and accelerated follow-up work enabled by prior investigation of the compounds resulted in identification of a series of pyrazolopyrimidine human phosphodiesterase (PDE)-V inhibitors with potent anticryptosporidial activity and efficacy following oral administration in *C. parvum*-infected immunocompromised mice. The novel PDE inhibitor leads (**compounds PDEi2** and **PDEi5**) affect parasite egress from infected host cells.

They have comparable activity against *C. parvum* and *C. hominis*, rapidly eliminate *C. parvum* in tissue culture, and have minimal off-target effects in a panel of safety screening assays. In comparison, the potent human PDE-V inhibitors sildenafil and the 4-aminoquinoline compound 7a have no useful activity against *C. parvum.* Based on homology modeling and *in silico* compound docking, **PDEi5** interacts directly with an active-site metal ion and docks well to two *C. parvum* PDEs. In contrast, larger amino acid side groups (Val900/Tyr11128 and His884/Asn1112) in both *C. parvum* PDEs replace alanine in human PDE-V and block sildenafil binding, explaining its lack of efficacy. These results identify a promising new drug target and lead series for anticryptosporidial drug development and validates a route to target-based optimization.

## Introduction

Cryptosporidiosis, the diarrheal disease caused by *Cryptosporidium* parasites, is one of the most important causes of life-threatening diarrhea in infants worldwide, causes incurable diarrhea in AIDS and transplant patients, and is the most identified cause of waterborne diarrheal outbreaks in the United States and Europe^1-3^. Cryptosporidiosis is also strongly associated with child malnutrition, growth stunting, and delayed cognitive development^4^. Two species, *Cryptosporidium hominis* and *Cryptosporidium parvum*, account for almost all human infections. Unfortunately, there is no protective vaccine, and the only approved treatment, nitazoxanide, is only modestly effective in children (∼56%) and equivalent to a placebo in AIDS patients^5,6^.

The need for improved anticryptosporidials has spurred research over the last ∼10 years, with various teams using phenotypic and/or target-based drug development approaches^7-20^. The current drug pipeline includes compounds targeting phosphatidyl-inositol-4-kinase (PI4K)^18^, calcium-dependent kinase 1 (CDPK1)^11-13^, cleavage and polyadenylation specific factor 3 (CPSF3)^19^, tRNA synthetase inhibitors (methionyl-, phenyl-, and lysyl-)^8,10,21^, and a partially optimized triazolopyridazine with unknown mechanism-of-action^20,22^. Two pre-clinical drug candidates are poised for first-in-human studies^18,19^. However, a recent dairy calf efficacy study of a methionine tRNA-synthetase inhibitor failed due to emergence of drug-resistant *C. parvum* with target gene mutations within days of beginning treatment^23^. Given the small number of validated targets and likelihood of drug resistance, there remains an urgent need to identify drug leads with novel mechanisms-of-action.

Since the heaviest burden of *Cryptosporidium* infection falls on young children in low- and middle-income countries, there is little financial incentive for anticryptosporidial drug development. The market for treatment of agricultural animals, especially cattle, is more favorable than for humans but still too small to stimulate de novo drug development. Realistic *Cryptosporidium* drug development efforts, as for any neglected disease, must therefore simultaneously respect the biological and pharmacological requirements of the intended use (i.e., for human cryptosporidiosis, treatment of small intestinal epithelial infection in young, often malnourished children and/or immunocompromised individuals) and the financial realities that impede target-based drug development. Drug repurposing has been widely touted as a solution to this mismatch between public health needs and financial reality^24^, and the approach, coupled with hypothesis, has successfully yielded new treatments for several conditions (e.g. remdesivir for Covid-19^25^). However, as has recently been highlighted regarding efforts to identify existing drugs for treatment of Covid-19, hypothesis-free screening for drug repurposing has resulted in no approved treatments despite widespread efforts and substantial research funding over approximately the last 20 years^26^. In the case of *Cryptosporidium*, numerous existing drugs inhibit parasite growth *in vitro* but lack activity in animal models^27^, possibly in part because anticryptosporidial drug efficacy appears to require sustained drug levels in the intestine while most clinically available drugs are optimized for oral absorption^28^. And clofazimine, a drug repurposing candidate with efficacy in one of two *Cryptosporidium* mouse models assessed, appears to be parasitistatic and failed in an early-stage human trial of AIDS patients with cryptosporidiosis^9,20,29^.

We believe a more promising approach than straight-forward drug repurposing is identification of chemical starting points by screening well-characterized compounds for which there has been substantial study for other purposes but that are at a development stage that permits lead optimization. This approach applies the principle of “Selective Optimization of Side Activities” (SOSA) to high quality, advanced compounds and leverages prior medicinal chemistry, pharmacokinetic (PK) and pharmacodynamic (PD) data, and safety studies to accelerate screening hit prioritization and early-stage optimization, while still enabling optimization of potency and PK/PD characteristics for specific uses^30^. Towards this end, a growing number of pharmaceutical companies are providing researchers in academia and other non-profits access to chemical libraries of well-characterized compounds.

Using this strategy, we now report whole-cell screening to identify *Cryptosporidium* growth inhibitors amongst well-characterized drug-like compounds made available through the Merck KGaA, Darmstadt, Germany Open Innovation Portal^31,32^. The included compounds possess a wide range of molecular targets encompassing enzymes (kinases, phosphatases, proteases, cyclic nucleotide phosphodiesterases, etc.), hormone or neurotransmitter receptors (serotonin, angiotensin II, endothelin, etc.), ion channels and several others. After confirmatory growth inhibition studies, anticryptosporidial screening hits were prioritized based on existing data and availability of analogs for early-stage optimization. The most promising compounds were then tested *in vivo*, which yielded a novel pyrazolopyrimidine phosphodiesterase-V (PDE- V) inhibitor series. The potent human PDE-V inhibitor sildenafil does not affect *C. parvum* growth, suggesting a target other than human PDE-V. Consistent with inhibition of *C. parvum* PDEs as the primary mechanism-of-action, the new lead series docks with high confidence to *C. parvum* PDE1 (CryptoDB cgd3_2320) and PDE2 (CryptoDB cgd6_4020), while sildenafil does not bind. This work collectively identifies the pyrazolopyrimidine PDE inhibitors as new high-value drug leads and establishes *Cryptosporidium* PDEs as a new drug target.

## Results

### Compound screening and down-selection of identified *C. parvum* growth inhibitors

We used a previously optimized high-content microscopy assay of *C. parvum* (Iowa) growth in the colon cancer cell line HCT-8^27^ to screen two compound libraries provided by the Merck KGaA, Darmstadt, Germany Open Innovation Portal, the Open Global Health Library and the Mini-Library (together, 278 unique compounds)^31,32^. Twenty three compounds selectively inhibited *Cryptosporidium* growth by over 70% at 5 μM (eight at 1 μM), yielding overall screening hit rates at 5 and 1 μM of 8.3% and 2.9%, respectively. Screening hits were selected for resupply and confirmatory dose-response testing based on availability and apparent potency, yielding thirteen confirmed inhibitors with 50% effective concentrations (EC50) ranging from 0.05 μM to 8.0 μM.

The confirmed *C. parvum* inhibitors were preliminarily prioritized based on a combination of potency and existing data regarding the mammalian target, PK characteristics, safety, and available chemical variants for follow-up testing. This yielded three chemical series of interest for further studies (Fig. 1): 1) PI3Ki1 (EC50 = 0.144 μM), a human phosphatidylinositol-3-kinase inhibitor; 2) PDEi1 (EC50 = 0.91 μM), a human phosphodiesterase V (PDE-V) inhibitor; and 3) IKK2i1 (EC50 = 4.1 μM), an inhibitor of the human inhibitor of NF- κB kinase 2 (IKK2). Of note, although the thiazolidine-2,4-dione side chain of PI3Ki1 is associated with nuisance compounds that give false positive screening results (termed pan-assay interference compounds (PAINS)) ^33^, existing data indicated no false positives with PI3Ki1 in other assays and published reports on related compounds indicated that they are highly selective and form non-covalent binding interactions with PI3K in crystal structures^34,35^.

**Figure 1:**
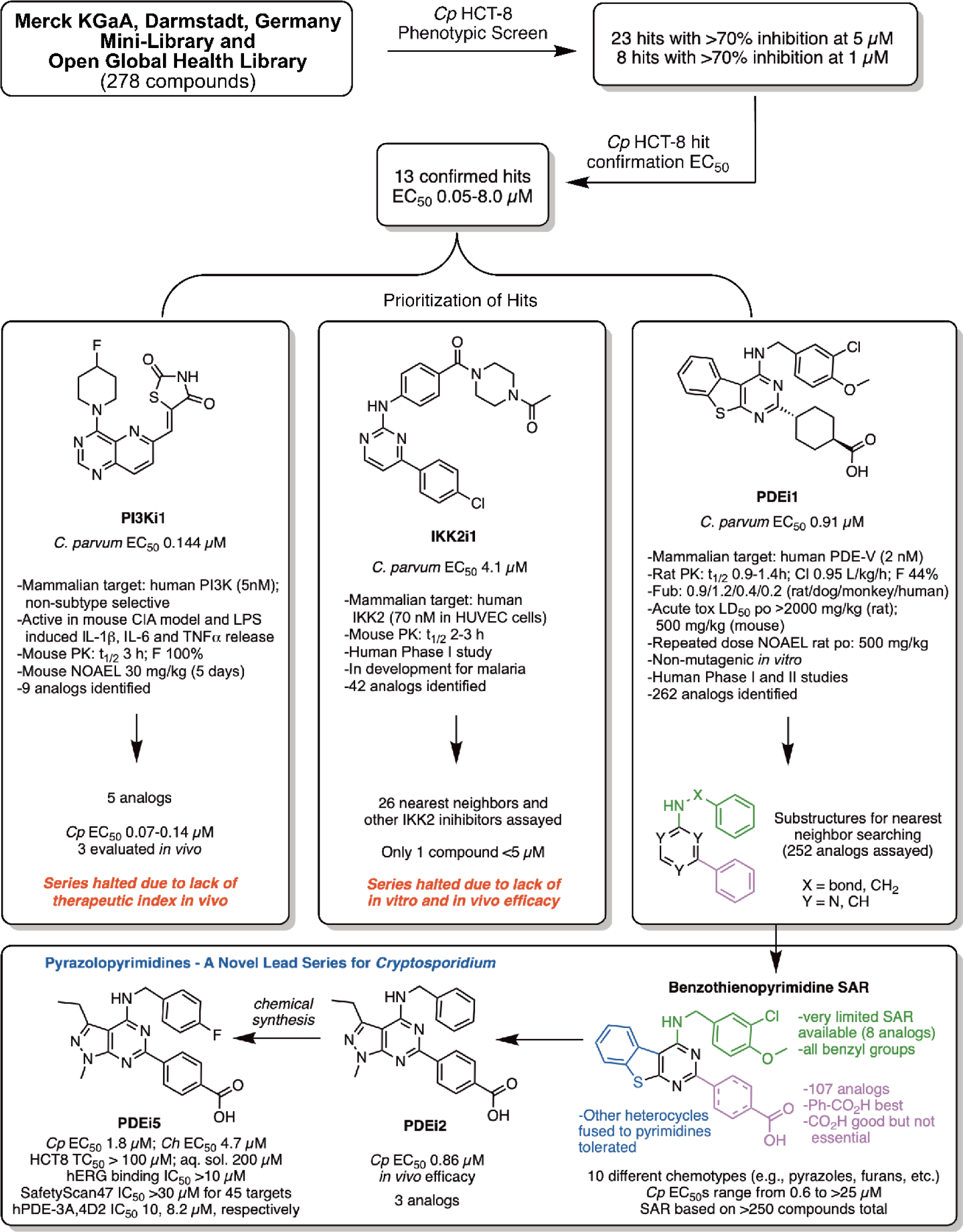
Screening results and prioritization considerations. The shown phosphatidyl-inositol-3-kinase inhibitor (PI3Ki), phosphodiesterase-5 inhibitor (PDEi) and IkappaB kinase 2 inhibitor (IKK2i) compounds were prioritized for follow-up based on confirmatory anticryptosporidial EC50 data, existing safety and pharmacologic data, and the availability of chemical analogs for initial follow-up studies.

## Nearest neighbor structure activity relationship development

We next conducted structure activity relationship (SAR) studies with analogs available in the chemical libraries provided by Merck KGaA, Darmstadt, Germany, both to judge the likelihood that significant progress might be made through a future medicinal chemistry program and to identify compounds worthy of testing for anticryptosporidial activity in animals. For this, the IKK2i, PI3Ki and PDEi chemical series were used for nearest neighbor (chemical similarity search^36^), chemical substructure, and target-based searches of the chemical libraries to identify compounds for dose-response testing. Of 26 IKK2i nearest neighbors and one other IKK2i series assayed, only one inhibited *C. parvum* at EC50 < 5 μM. The lone exception was IKK2i2, an unrelated pyrazole chemotype IKK2 inhibitor (Table S2) that was exceptionally potent with an EC50 of 0.020 µM. Only five total PI3Ki analogs were available for follow up testing that had *C. parvum* EC50s ranging from 0.070 to 0.140 μM.

On the other hand, 252 PDEi1 benzothienylpyrimidine analogs and an additional ten closely related chemotypes were available for follow up testing (Fig. 1; Table S2). *C. parvum* EC50s for these series ranged from 0.6 to >25 μM. Most of the SAR was provided by the benzothienopyrimidines. Limited data was available for the *N*-benzyl position (green in Fig. 1) as only eight comparator analogs were available and all were *N*-benzylated. A much richer SAR was gleaned from the cyclohexyl/benzoic acid position (magenta in Fig. 1) where benzoic acid (e.g., PDEi3) appeared to be the preferred substituent and COOH was preferred off the aryl ring but was not essential. Finally replacement of the benzothiophene was tolerated by other chemotypes (blue in Fig. 1) such as benzofurans, pyrazoles (e.g., PDEi2), indoles (e.g., PDEi4), thiophenes and pyrroles. From this analysis, 1-methyl-3-ethylpyrazolopyrimidines were a preferred chemotype relative to the others and comparable in potency to the benzothienopyrimidines while providing potential for improved pharmacokinetics due to reduced sp^2^ character. These three pyrazolopyrimidines had potency comparable to the original hit and differed only in substitution off the benzyl ring (*N*-benzyl, *N*-CH2(4-chlorophenyl) (PDEi2), and *N*-CH2(4-methoxy-3-chlorophenyl)). These three analogs all had *C. parvum* EC50s below 1 µM.

In an effort to see if the early SAR trend held, we designed a simple 4-fluorobenzyl analog of PDEi2, rationalizing that if 4-H and 4-Cl were potent, the 4-F would also be potent and make a useful comparator. This *N*-CH2(4-fluorophenyl) (PDEi5) analog was synthesized over 5 steps from a known aminocyanopyrazole intermediate **I** (Supplemental Fig. 1)^37^. Nitrile **I** was hydrolyzed, condensed with urea, and chlorinated with PCl3/PCl5 to give dichloropyrimidine intermediate **IV**. Selective nucleophilic displacement with 4-fluorobenzylamine followed by Suzuki coupling and ester hydrolysis gave compound PDEi5. PDEi5 was found to have similar potency with *C. parvum* EC50 of 1.8 µM.

### *In vivo* efficacy and pharmacokinetics

*Cryptosporidium parvum* infected NOD SCID gamma (NSG) mice were used to assess efficacy of the PI3Ki, IKK2i, and PDEi series *in vivo*^20^. Using existing *in vitro* metabolic stability and mouse PK data on the chemical analogs studied, nine compounds (two IKK2i, three PI3Ki, and four PDEi) were selected and tested for *in vivo* efficacy (Table S1 shows characteristics of compounds tested *in vivo* and Table S2 provides compound structures). Both IKK2is tested were ineffective, and all three PI3Kis were toxic at the dose tested, necessitating study cessation after just two days of dosing due to weight loss. The most promising PI3Ki (PI3Ki1) was retested at a dose of 10 mpk twice daily, below NOAEL of 30 mg/kg known from previous analyses, and proved to be ineffective. On the other hand, three of four PDEis tested in the proof-of-principle experiment appeared effective (p < 0.05) or partially effective with a 4-day dosing and no adverse effects were noted (Supplemental Fig. 2).

The pyrazolopyrimidines PDEi2 and PDEi5, and the indolopyrimidine PDEi4 were studied in follow-up with 7-day dosing regimens in two independent experiments (Fig. 2). PDEi4 had no significant effect. PDEi2 (Fig. 2A) and PDEi5 (Fig. 2B), in contrast, both reduced parasite shedding by ∼99% after 7 days of treatment (p = 0.02), although infection relapsed within a week of stopping treatment.

**Figure 2:**
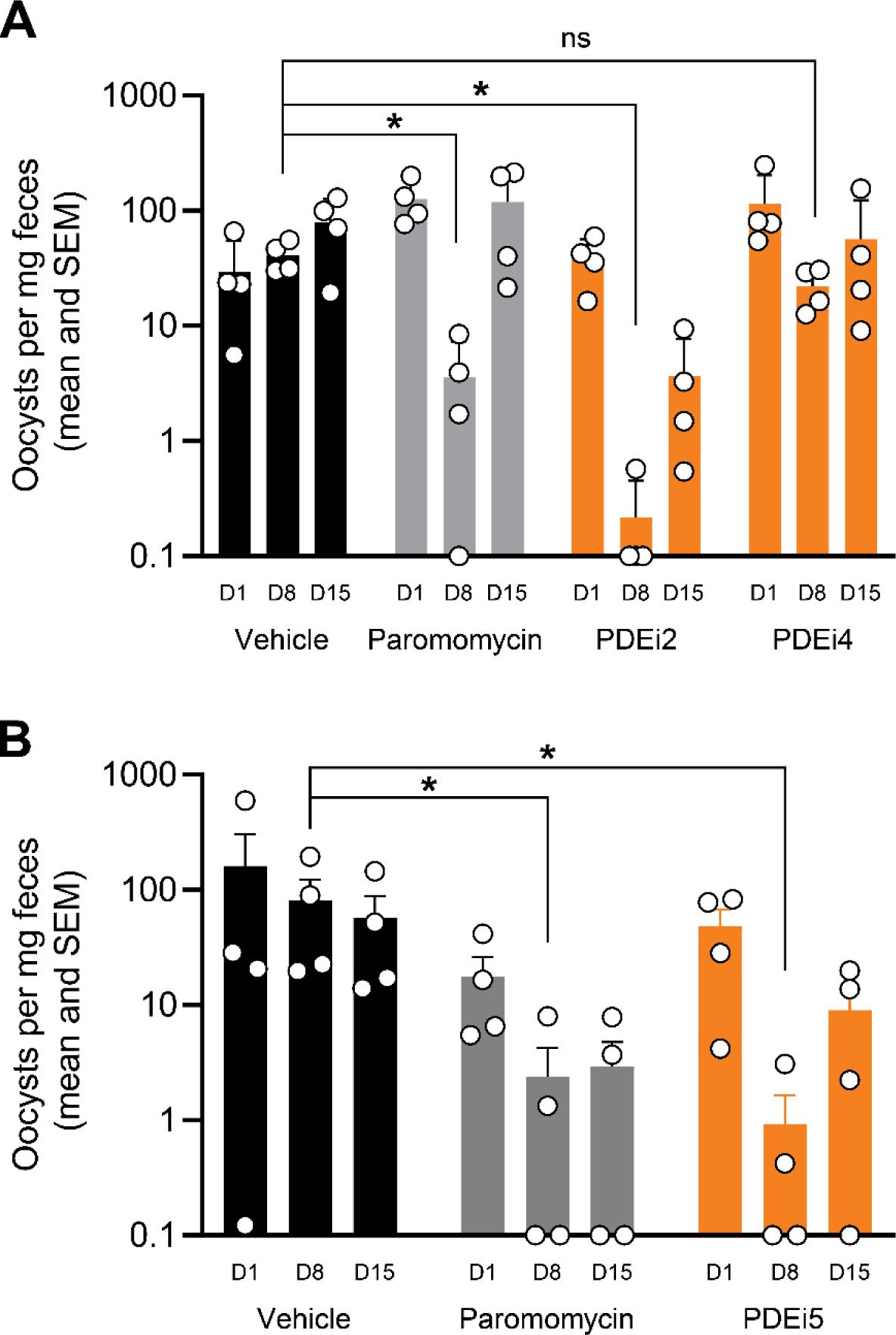
PDEi efficacy in established murine *C. parvum* infection. The indicated compounds were tested for efficacy in established cryptosporidiosis in NSG mice. *C. parvum* infection was allowed to progress for either 7 days **(A)** or 14 days **(B)** prior to oral administration of 50 mpk twice daily of the indicated compounds or 1000 mpk twice daily of paromomycin (positive control) by oral gavage for 7 days. Parasite shedding in the feces was determined at the onset of treatment (D1), completion of treatment (D8), and one week after completion of treatment (D15). Data are the mean and SEM (n = 4 mice per experimental group) of parasite fecal shedding per mg of feces, and data points below the limit of qPCR detection are shown on the x-axis. Asterisk (*) indicates p > 0.05 vs. vehicle control for the indicated experiment by Kruskal-Wallis test. ns indicates not significant.

PDEi2 had kinetic solubility = 190 μM (2h, PBS, pH 7.4) and other favorable properties for *in vivo* work (see Table S1). Single dose IV and oral mouse PK studies were therefore performed with PDEi2 to further understand the determinants of *in vivo* efficacy for this series. PDEi2 was rapidly cleared from plasma (T1/2 ∼ 1 h) following IV administration (Fig. 3A). In contrast, after a single 10 mg/kg oral dose, total drug concentrations in successive intestinal tissue segments persisted at high levels for at least 7 hours and were still detectable after 24 hours (Fig. 3B). Because tissue lysates mix interstitial and intracellular fluids, we cannot know how drug is partitioned between the extracellular and intracellular compartments ^38^. We also cannot accurately determine the free (unbound) concentration in the lysate, since its protein composition is different from plasma. Thus, the precise unbound concentration at the site of action, presumably within the parasite which itself is located within the parasitophorous vacuole in the enterocyte, cannot be readily determined. Nevertheless, these data show that the total PDEi2 concentration in the intestinal tissue persisted for hours at levels that exceeded multiples of the EC90 for inhibition of parasite growth, which is consistent with observed parasiticidal activity in *in vitro* time-kill assays (see Fig. 4). Drug concentrations in plasma were also prolonged with oral versus IV administration, likely due to continued gastrointestinal absorption, but did not reach the same levels measured in intestinal tissues (Fig. 3A).

**Figure 3:**
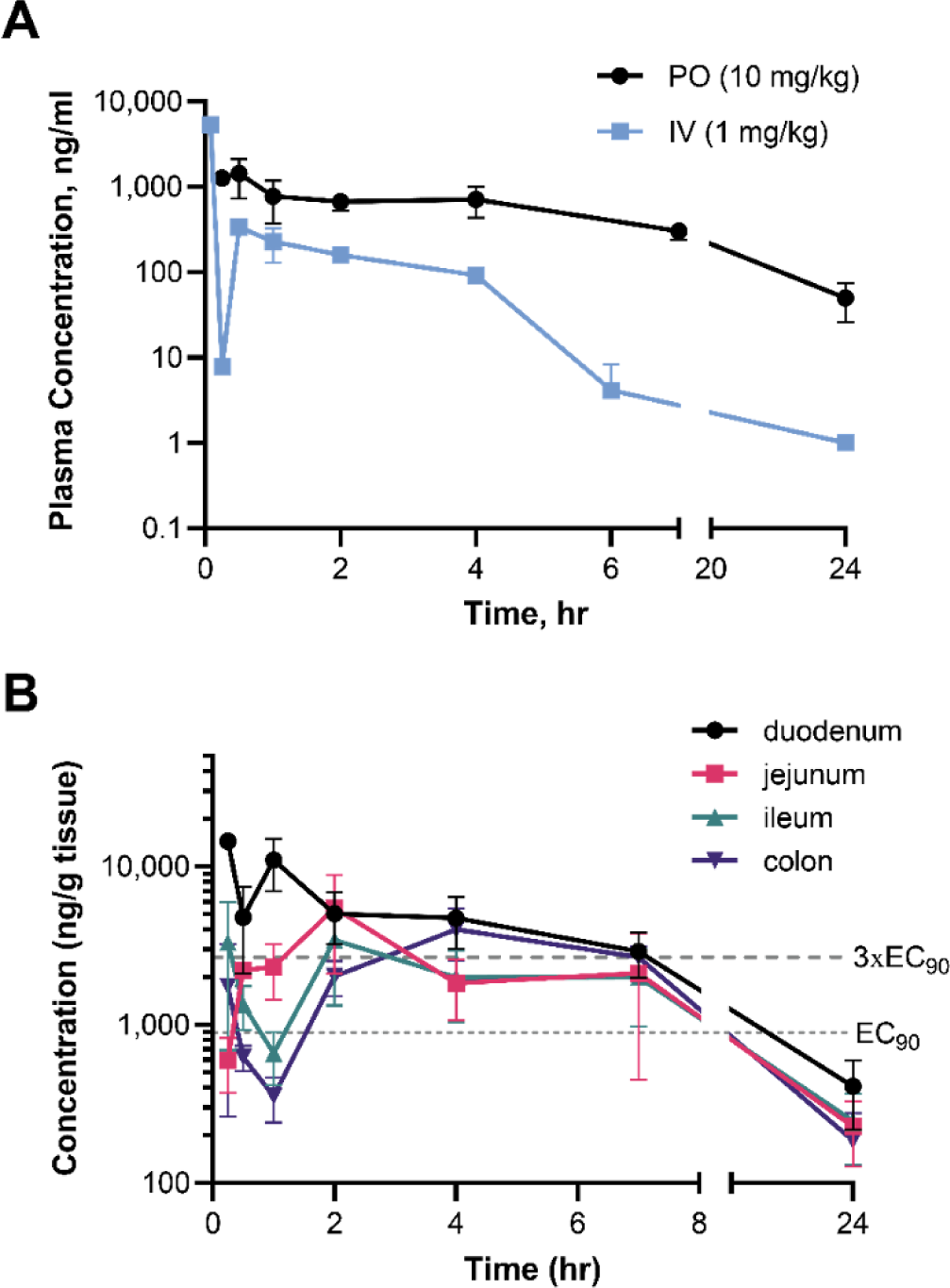
Mouse PK data for compound PDEi2. (A) Time courses of total drug concentration determined in plasma with single dose IV and oral administration at indicated doses. **(B)** Time courses of total drug concentration determined in washed intestinal tissue segments with single dose 10 mg/kg oral administration. Data for (A) and (B) are mean and SE (n=3 mice per experimental group).

**Figure 4:**
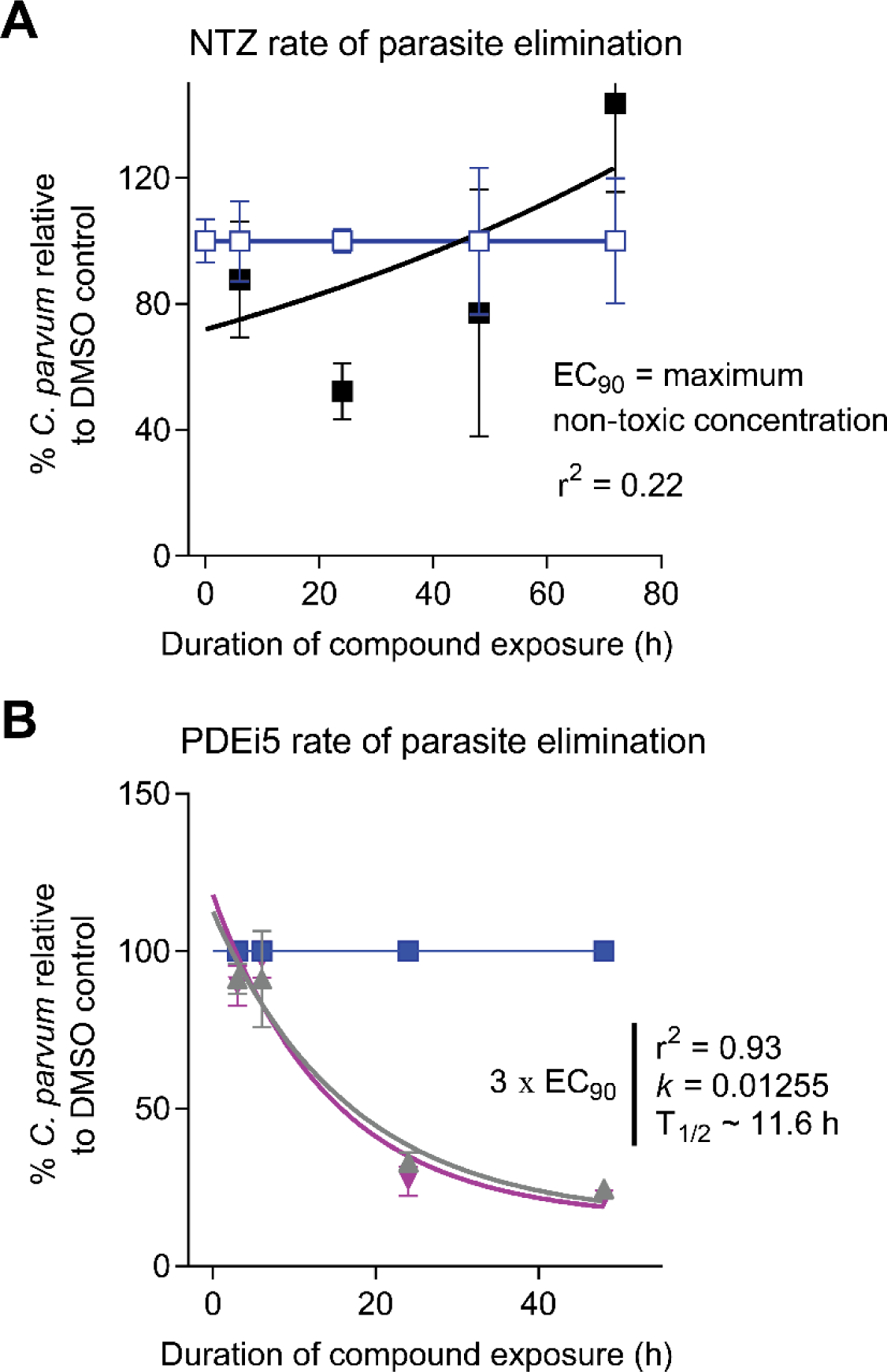
I*n vitro* rate of parasite elimination. Time-kill curves for **(A)** nitazoxanide (NTZ) and **(B)** PDEi5 showing elimination of *C. parvum* from infected HCT-8 cells. One-phase exponential decay curves were fit for NTZ and PDEi5. Data are the means and SD for 4 culture wells per time point, normalized for each time point to the vehicle control (DMSO). **(A)** No exponential decay curve could be fit for NTZ at the EC90 which was the highest non-toxic concentration. Blue (DMSO); black (NTZ EC90). **(B)** Rapid parasite elimination by PDEi5. Curves are shown for 3×EC90 (grey) and 6×EC90 (cyan)(the highest concentration tested), or the DMSO control (blue).

### Activity against *C. hominis*, rate-of-action, and anticryptosporidial mechanism-of-action

Most human cryptosporidiosis is caused by either *C. parvum* or *C. hominis*, but most work (including our screening assays and confirmatory studies) is done using *C. parvum* because *C. hominis* oocysts can only be sourced from patients or using an expensive gnotobiotic piglet model. Nonetheless, confirmation of activity against both species is critical prior to investing time and resources into a given drug lead series. We used the *C. hominis* Tu502 strain and the cell-based assay method used for *C. parvum* to test the activity of PDEi2, PDEi4, and PDEi5. Activity was comparable for both species (see Table S1); for new lead compounds PDEi2 and PDEi5, the *C. hominis* EC50s and 95% confidence intervals were 0.61 (undetermined-3.76) μM and 4.7 (2.4-9.1), respectively.

Given the desire to develop anticryptosporidials with efficacy for immunocompromised patient populations, we use time-kill curves at early stages of our developmental cascade to enable prioritizing compounds that rapidly eliminate *C. parvum* in tissue culture and are likely parasiticidal^20^. Figure 4 shows time-kill data in the HCT-8 cell culture system for nitazoxanide and PDEi5. Decay curves were fit after referencing data at each time point to the vehicle control to isolate the effect of compounds from the spontaneous decline in *C. parvum* numbers that begins after 48-60 hours of *in vitro* culture. Like previously reported data suggesting parasitistatic activity^19,20^, no nitazoxanide-dependent parasite elimination occurred at the highest tolerated dose. PDEi5, on the other hand, drove progressive parasite elimination with an exponential decay half-life of ∼ 11.6 hours. Thus, approximately 116 hours (<5 days) would be required for 99.9% parasite reduction by PDEi5 in the absence of host immune help. Consistent with a class effect which has been typical with this time-kill assay, results were similar with PDEi2 (Supplemental Fig. 3). These data were also in keeping with the *in vivo* efficacy in the NSG mouse with twice daily administration, since the intestinal tissue concentration of PDEi2 following a single 10 mg/kg dose exceeded the EC90 for over 7 hours.

PDEi2 potently inhibits the mammalian cGMP-specific phosphodiesterase PDE-V (IC50 = 20 nM) and less potently inhibits other cGMP-specific phosphodiesterases, including PDE-I, PDE-II, PDE-IV, and PDE-VI (Table S3). To determine if the anticryptosporidial activity of the PDEi series resulted from inhibition of host PDE-V or parasite phosphodiesterases, we compared the anticryptosporidial activity of two PDEi analogs (PDEi1 and PDEi5; structures given in Table S2) to the potent mammalian PDE-V inhibitors sildenafil and the 4-aminoquinoline compound 7a (Sigma-Aldrich, cat#508957) (IC50 for human PDE-V = 3.5 nM and 0.28 nM, respectively)^39,40^. The EC50 of sildenafil for *C. parvum* growth in HCT-8 cells exceeded 50 μM and that of compound 7a was 36 μM, far in excess of the concentrations needed to inhibit host PDE-V and suggesting that *Cryptosporidium* growth inhibition by the new PDEi series was independent of host cell PDE-V and more likely resulted from inhibition of one or more *Cryptosporidium* phosphodiesterases or another off-target effect (Supplemental Fig. 4A).

We next used available AlphaFold structure models^41,42^ for the *C. parvum* PDEs and X-ray crystal structures of human PDE-V (*h*PDE5) to determine if the new PDEi series compounds are likely to bind *C. parvum* PDEs and to understand the inability of sildenafil to block *Cryptosporidium* growth. Alpha-fold models were available for two of the three predicted *Cp*PDEs identified by BLAST of *h*PDE5A (sp|076074) against the *Cryptosporidium* genome (CryptoDB: cgd3_2320 (E=9e-18) and cgd6_4020 (E=1e-08), henceforth *Cp*PDE1 and *Cp*PDE2, respectively), although neither protein has been characterized experimentally. Published transcriptomics data indicate *Cp*PDE1 is highly expressed in sporozoites, asexual forms, and less conclusively, in female gametocytes. *Cp*PDE2, on the other hand, is minimally expressed in sporozoites and asexual forms, and its expression increases dramatically at 48 hours in HCT-8 cell cultures when gametocytes become predominant^43^. Only *Cp*PDE1 is predicted to be cGMP-specific like *h*PDE5A, with homology to *h*PDE5A limited to 260 of 997 total amino acids (29% identity, 46% similarity) that correspond to the predicted *Cp*PDE1 active site (sequence alignment in Supplemental Fig 5A). The existing AlphaFold *Cp*PDE models (AlphaFold: A3FQ29-F1-model_v4 (*Cp*PDE1) and A3FPW6-F1-model_v4 (*Cp*PDE2)) have high confidence predicted structures starting near GLY590 and ASP825, respectively. This high confidence region corresponds to the *C*-terminal catalytic domain. However, the AlphaFold structures lacked active site metal ions known to interact with cAMP/AMP that would potentially interact with the novel PDE inhibitors. Therefore, to enable addition of the active site Zn^2+^ and Mg^2+^ metal ions to the models, we overlayed the crystal structure of a *Schistosoma manosoni* PDE (*Sm*PDE4 (pdb 6EZU^44^)) that is 36% identical and 53% similar to *Cp*PDE1 in the active site. The AlphaFold models were truncated to remove the low-confidence and non-catalytic *N*-terminal region, and the active site metal ions and waters from the *Sm*PDE structure, also observed in related PDEs, were incorporated by merging followed by the restrained minimization routine using the OPLS4 forcefield in Schrödinger Maestro and aligned to produce structures suitable for molecular docking studies (Supplemental Fig. 5B). Using these models, PDEi5 was docked to both *Cp*PDE1 and *Cp*PDE2 using the SP Glide docking routine in Schrödinger Maestro^45^. Docking scores for the top three poses in both models ranged from -9.5 to -9.9 and -10.3 to -10.8 for *Cp*PDE1 and *Cp*PDE2, respectively, indicative of strong binding interactions. The models were further refined using the Induced Fit docking routine in Schrödinger Maestro, giving docking scores as low as -11.9 and -11.7 for *Cp*PDE1 and *Cp*PDE2, respectively, suggesting PDEi5 has a high likelihood of binding these targets (Fig. 5 and Supplemental Fig. 6). All of the docked poses of PDEi5 in both structures involved a direct interaction of the carboxylic acid moiety with the two metal ions, mimicking the cGMP/cAMP phosphate interaction with the metals observed in *Sm*PDE4. While positioning of the benzyl and pyrazolopyrimidine core varied, structurally conserved Phe935/1161 formed π-π stacking with PDEi5. Both PDEi5 and sildenafil docked with similar docking scores in human PDE-V, but sildenafil docked poorly to the *Cp*PDE models (best docking scores were -5.9 and -3.6, respectively) due to its inability to adopt the same binding pose it has in the human structure, likely due to the bulky 2-ethoxy-5-sulfonamide-aryl ring (Supplemental Fig. 7). Ala873 in *h*PDE-V is replaced by more bulky Val900 and Tyr1128 in *Cp*PDE1 and *Cp*PDE2, leading to a clash with the ethoxy group of sildenafil. Furthermore, Ala767 in *h*PDE-V is replaced by more bulky His884 and Asn1112 in *Cp*PDE1 and *Cp*PDE2, leading to clashing with the *N*-Me of sildenafil. Of note, seven of the 18 residues within 4Å of the PDEi5 docked to *Cp*PDE1 differ from *h*PDE-V, suggesting potential to design variants of PDEi5 with improved selectivity for *Cp*PDE1 over *h*PDE-V.

**Figure 5:**
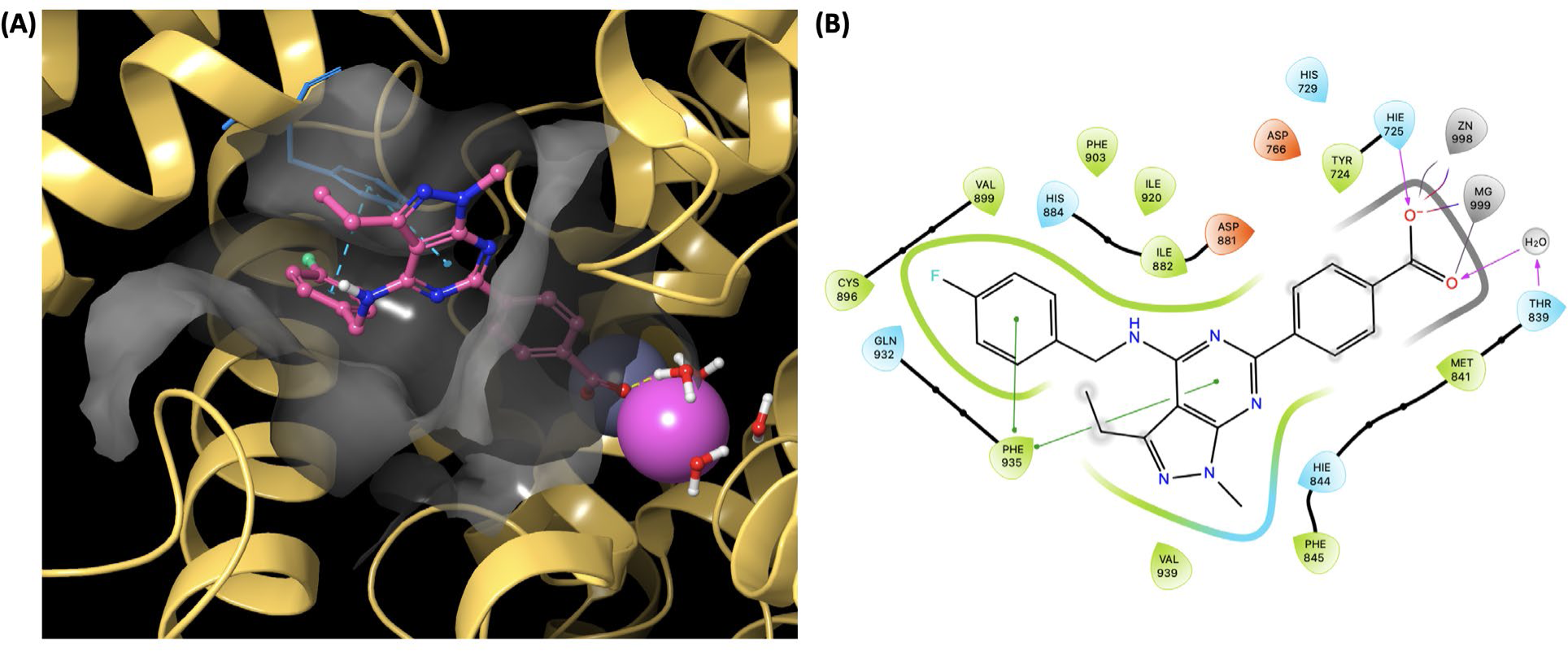
Lowest energy pose from induced fit docking of PDEi5 in the *Cp*PDE1 model. Glide docking score -11.9. The ligand binding pocket of *Cp*PDE1 is shown as a gray surface. (A) Binding site illustrating Phe935 (blue) forming π-π face-face and edge-face interactions with PDEi5. (B) Ligand interaction diagram illustrating residues within 4Å of the ligand, π-stacking between ligand and Phe935 (green lines), and ionic salt bridge and hydrogen bonding between the carboxylate of the ligand with the zinc, magnesium, His725 and water molecules.

Finally, we employed a suite of *C. parvum* life cycle-based assays^46^ to begin investigating the mode-of-action of the new PDEi series, and the likely function(s) of *Cp*PDEs in *Cryptosporidium* biology. The PDEi inhibitors were inactive in a cell invasion assay (Supplemental Fig. 4B) but strongly active in an assay that measures a combination of *Cryptosporidium* egress and invasion of new host cells (Supplemental Fig. 4C), suggesting their primary activity is to block parasite egress. They also blocked sexual maturation, since they reduced expression of the macrogametocyte marker DMC1 (Supplemental Fig. 4D). Since anticryptosporidial pipeline diversity is critical and the mechanism of many compounds in development is unknown, we used a clustering algorithm previously reported with these phenotypic assays and publicly available assay data to generate a dendrogram and assess if the PDEi’s mode-of-action was distinct (Supplemental Fig. 8)^46^. Both PDEis clustered together, as expected, and separately from other compound series in development.

### Safety assessment and identification of potential liabilities

Pyrazolopyrimidine PDEi2 was non-toxic to the HCT-8 cell line at 100 μM, the highest concentration tested (note that per our data, the kinetic solubility of PDEi2 is 190 μM); selectivity of lead PDEi2 exceeded 125-fold. The 50% toxic concentration (TC50) of the indolopyrimidine PDEi4 (kinetic solubility 21 μM) was 87 μM (selectivity index = 100). This in combination with superior activity in the mouse led us to focus further safety assessment on the pyrazolopyrimidine series.

We contracted Eurofins (St. Charles, MO) to conduct off-target safety profiling for PDEi5, the pyrazolopyrimidine PDEi2 analog that we had synthesized independently in our lab. PDEi5 was tested in dose-response up to 30 μM in the SafetyScan47 panel of cell-based assays that assesses effects on 47 human targets (including *h*ERG; 78 assays total, as many of the targets are run in both agonist and antagonist modes)^47^. PDEi5 had EC50 values >30 μM for 45 of 47 targets (see Table S4 for full list of targets and assay results). The only off-target enzymes inhibited were human PDE-IIIA and PDE-IVD2 with EC50 values of 10.2 and 8.2 μM, respectively. To assess human PDEs not included in the SafetyScan47 panel, we contracted with Eurofins to measure PDEi5 inhibition of 13 mammalian PDEs (Table S5). PDEi5 most potently inhibited human PDE-V and PDE-VI, but PDE-IVB1 and PDE-XIA were also inhibited >50% at a test concentration of 10 μM. PDEi5 was also tested by Eurofins to demonstrate it has suitable aqueous solubility (200 µM, kinetic) and has low potential to inhibit the hERG channel (-14.7% at 10 µM in a [^3^H]-dofetilide competitive binding experiment). We also evaluated the potential for toxicity of PDEi5 using *in silico* methodology using ProTox-II, which indicated a low probability for toxicity (Supplemental Fig. 9)^48^. Finally, Eurofins tested *in vitro* inhibition of a panel of seven cytochrome p450s (CYPs), demonstrating modest inhibition of CYP2C8 and CYP2C9 (∼31% and 18% respectively at 10 μM) (Table S6).

## Discussion

This work establishes *Cryptosporidium* PDEs as new targets for anticryptosporidial development and the pyrazolopyrimidine phosphodiesterase inhibitors (i.e., PDEi2 and PDEi5) as a promising drug lead series. New leads PDEi2 and PDEi5 possess many desirable attributes as starting points to develop an improved *Cryptosporidium* treatment, since they rapidly eliminate parasites *in vitro*, have comparable activity against both *C. parvum* and *C. hominis*, are highly efficacious in a stringent mouse model of infection, have minimal off-target effects on a panel of human targets tested, and have only modest potential for drug-drug interactions due to CYP inhibition. Homology/docking studies done using high-quality AlphaFold structures predict that the new PDEis bind well to *Cp*PDE1 and 2, providing a likely mechanism-of-action. Steric clashing predicted to reduce binding affinity of sildenafil to *Cp*PDEs likely explains its inability to inhibit *Cryptosporidium* growth, further supporting that the new PDEi series affects *Cryptosporidium* by targeting the parasite enzyme rather than host PDEs. Phenotypic assays with the compounds suggest a primary function of *Cp*PDE1 in the HCT-8 cell culture system (note that *Cp*PDE2 is minimally expressed during the relevant growth phases) in parasite egress from host cells. Finally, the overall pattern of effects in phenotypic assays indicates that the new pyrazolopyrimidine PDEi series works via a mechanism that is distinct amongst current compound series in the developmental pipeline, and thus contributes important drug pipeline diversity. These studies collectively establish the pyrazolopyrimidines as a high priority series for further development and suggest the possibility of target-based optimization.

It is likely that nitazoxanide fails in malnourished children and immunocompromised people because it is parasitistatic rather than parasiticidal for *Cryptosporidium* species and requires assistance from the host immune system to clear cryptosporidiosis; unlike nitazoxanide, PDEi2 and PDEi5 rapidly eliminate *C. parvum* from *in vitro* culture. A growing body of published data indicate that *in vivo* efficacy in immunocompromised animals correlates with a compound’s rate-of-action^10,19-21^; furthermore, rapid vs. slow or static anticryptosporidial activity appears to be conserved amongst chemical analogs with a given mechanism of action. Since PDEi2 and PDEi5 rapidly eliminate *C. parvum* from tissue culture, we believe the pyrazolopyrimidines are a lead series for which optimization could yield a treatment effective for all patient groups impacted by cryptosporidiosis.

Consistent with this, PDEi2 and PDEi5 are efficacious in an NSG mouse model that tests the ability to impact established infection. Numerous mouse models have been used for anticryptosporidial efficacy studies, but most mice infected by *C. parvum* develop an acute infection that resolves spontaneously over approximately 17 days^9,49-51^. In the absence of effective drugs to benchmark different models, the utility of any animal model for predicting efficacy in children and immunocompromised people with cryptosporidiosis is, of course, unknown. However, we use the NSG mouse model, because NSG mice develop persistent infection localized to the ileum and cecum that enables delaying study drug administration until infection is fully established and monitoring for recrudescence after treatment cessation^20^. Relative to self-resolving infection models, the NSG mouse model of cryptosporidiosis is stringent, and compounds active in acute models of infection are frequently inactive in the NSG mouse model (e.g., clofazimine^9^). PDEi2 is rapidly cleared from the plasma but persists in intestinal tissue at levels above the EC90 for parasite growth inhibition. Consistent with a growing body of data for other anticryptosporidial lead series, therefore, the key determinant of *in vivo* efficacy of the PDEi series appears to be adequate compound presence in the intestinal lumen and tissue^28^. In the case of PDEi2 and PDEi5, the maximal rate of parasite elimination *in vitro* was achieved at a concentration of 3x EC90, which provides a target tissue concentration to guide drug optimization.

Safety is essential for drugs intended for outpatient administration to young children in whom drug toxicities are difficult to monitor. PDEi2 is non-toxic to HCT-8 cells at concentrations exceeding 100 μM, yielding a selectivity index for *C. parvum* greater than 125. Furthermore, testing close analog PDEi5 with the Eurofins SafetyScan47 assay panel to screen for off-target effects only identified modest potency for inhibition of several additional mammalian PDEs, and *in silico* safety evaluation of PDEi5 using ProTox-II also indicates a low probability of toxicity^48^. Finally, minimal CYP450 inhibition suggests a low likelihood of drug-drug interactions that might interfere with treating AIDS patients or patients on immunosuppressive drugs following organ transplantation.

Based on structural modeling/docking studies in combination with phenotypic assays, the new PDEi series appears to impact *Cryptosporidium* growth by inhibiting at least one of three predicted *Cryptosporidium* PDEs which affects parasite egress from infected host cells, an essential step in progression of the asexual parasite life cycle. PDEi5 docks well to both modeled *Cp*PDE structures. Based on expression of *Cp*PDE1 in asexual stages and predominance of *Cp*PDE2 in sexual stages, it is likely that the effect of pyrazolopyrimidines on *Cryptosporidium* growth results from *Cp*PDE1 inhibition, but inhibition of *Cp*PDE2 may be important for *in vivo* efficacy. Genetic studies using CRISPR/Cas-9 to over-express or express mutant enzymes resistant to pyrazolopyrimidine inhibition will be needed to formally prove that the anticryptosporidial effect of the new series is due to *Cp*PDE inhibition and if *in vivo* efficacy requires inhibition of both enzymes. Such information could be used in combination with *in vitro* assays using recombinant enzymes to guide drug optimization. Furthermore, evaluation of structural models of PDEi binding suggests the potential for improving selectivity for the *Cryptosporidium* PDEs over human PDE-V through medicinal chemistry optimization due to 39% variation in active site residues (*Cp*PDE1 vs *h*PDE-V). In support of such optimization strategies, we are currently working to express functional recombinant *Cp*PDEs and make transgenic parasites engineered to overexpress the different *Cp*PDEs.

Despite the critical role of phosphodiesterases in controlling levels of cellular cAMP and cGMP and their proven druggability, there is limited precedence for targeting parasite phosphodiesterases for drug development. *Trypanosoma brucei* PDEB1 is a chemically validated therapeutic target for Human African Trypanosomiasis (HAT) for which the most advanced compounds are curative in a mouse model^52-56^. Benzoxaborole and other inhibitors of *Schistosoma mansoni* PDE4A (the enzyme used here to incorporate metal ions into the *Cp*PDE1 and *Cp*PDE2 AlphaFold models) have been identified^57,58^, but the best compounds only modestly reduce worm burden in *S. mansoni* infected mice, raising questions about suitability of the target ^59,60^. For Apicomplexa like *Cryptosporidium*, potent inhibitors of *Plasmodium falciparum* and *Toxoplasma* phosphodiesterases have been identified and proven useful to demonstrate the function of cyclic nucleotides in Apicomplexa^61^. Consistent with the effects of PDEi1 and PDEi5 on *Cryptosporidium* cell egress and reinvasion, these inhibitors appear to block *Plasmodium* proliferation by disrupting protein kinase G-dependent cell egress^61^. A medicinal chemistry effort has produced more potent inhibitors of *Plasmodium* PDEs, but *in vivo* efficacy has not been reported^62^. Due to their different locations in the host, the drug pharmacodynamics required for treating malaria, babesiosis, toxoplasmosis, and cryptosporidiosis differ and make a pan-apicomplexan drug unlikely. In addition to anticryptosporidial lead optimization studies, however, pathogen-hopping studies are warranted to determine if the new pyrazolopyrimidine anticryptosporidial drug leads described here can provide starting points to develop drugs for other Apicomplexa.

## Materials and Methods

### Chemical compounds

The Merck KGaA, Darmstadt, Germany Mini-Library and Open Global Health Library were supplied through the Open Innovation Portal as 10 mM DMSO stock solutions in microtiter plates. All screening hits available for follow-up were reacquired for confirmatory dose-response assays. Nearest neighbor and chemical substructure searches were performed using available Merck KGaA, Darmstadt, Germany compound libraries to further prioritize the confirmed screening hits for follow-up and available compounds were sourced directly from Merck KGaA, Darmstadt, Germany. The PDEi analog PDEi5 was synthesized by the Meyers Lab (St. Louis University); compound structures and purity ≥95% were verified by ^1^H NMR and LC-MS. Sildenafil (Sigma-Aldrich, cat#SML3033) and compound 7a (Sigma-Aldrich, cat#508957) were purchased for use in phenotypic assays.

#### General Synthesis Procedure

Unless otherwise stated, all reagents and solvents purchased were used as received without further purifications. Normal and reverse-phase chromatography were performed using CombiFlash® Rf+ (Teledyne Isco) with SiliaFlash F60 40−63 μm (230−400 mesh) silica gel (SiliCycle Inc.) eluting with hexanes/ethyl acetate gradient and RediSep Rf Gold pre-packed C18 cartridge or prep-HPLC on a ACCQ-Prep HP150 (RediSep Prep C18, 100Å, 5 µm, 250 mm x 20 mm or 30 mm column) eluting with acetonitrile/water gradient. Thin layer chromatography was carried out using TLC Silica gel 60 F254 Glass plates 2.5 x 7.5 cm (Merck KGaA, Darmstadt, Germany) and visualized with a UV lamp (254/365nm UV/6-watt, BioGrow®). Liquid chromatography mass spectrometry (LCMS) was performed with an Agilent 1100/1946 HPLC/MSD electrospray mass spectrometer in positive ion mode with a scan range of 100−1000 Da. ^1^H NMR spectra of intermediates and final compound were recorded and acquired in CDCl3 or DMSO-*d*6 as solvents using Bruker 400 MHz spectrometer at ambient temperature (400 MHz for ^1^H). Chemical shifts for ^1^H NMR (400 MHz) spectra are reported in parts per million (ppm) from either CDCl3 (7.26 ppm) or DMSO-*d*6 (2.50 ppm) with multiplicity (s = singlet, bs = broad singlet, d = doublet, t = triplet, q = quartet, dd =doublet of a doublet, td = triplet of a doublet, and m = multiplet) and coupling constants (*J*) in Hz. High resolution mass spectrum (HRMS) was obtained with an ABSciex 5600+ instrument. The verified purity of final compound was ≥95% as determined by HPLC UV absorbance unless noted otherwise.

#### 5-amino-3-ethyl-1-methyl-1*H*-pyrazole-4-carboxamide (II)

Concentrated sulfuric acid (H2SO4, 35 mL) was poured into a round-bottom flask and cooled in an ice water bath to 0 °C. 5- Amino-3-ethyl-1-methyl-1*H*-pyrazole-4-carbonitrile (**I**, 8.12 g, 54.1 mmol) was slowly added to the cooled concentrated H2SO4 at 0 °C ^37^. The ice bath was removed, and the resulting solution allowed to warm to room temperature with stirring overnight. After this time, the reaction was poured on an ice and neutralized with concentrated sodium hydroxide solution to pH 8. The aqueous was transferred into a separatory funnel and extracted 3x with ethyl acetate. The organics were combined, washed with brine, and dried over magnesium sulfate before filtration. The filtrate was concentrated on a rotary evaporator and the resulting light brown solid product (5.1 g) was > 98% purity on LCMS with a yield of 56% and was used in the subsequent step without further purification. ^1^H NMR (400 MHz, DMSO-*d*6) δ 6.50 (br. s., 2H), 6.13 (s, 2H), 3.46 (s, 3H), 2.65 (q, *J* = 7.46 Hz, 2H), 1.13 (t, *J* = 7.46 Hz, 3H). LC-MS m/z (M + H)^+^ = 169.

#### 3-ethyl-1-methyl-1*H*-pyrazolo[3,4-*d*]pyrimidine-4,6(5*H*,7*H*)-dione (III)

Urea (6.35 g, 106 mmol) and 5-amino-3-ethyl-1-methyl-1*H*-pyrazole-4-carboxamide (**II**, 2.60 g, 15.5 mmol) were weighed into a round bottom flask. The flask was heated to 200 °C for 2 h. The reaction was cooled to room temperature and 2M NaOH solution was added to the flask. The resulting solution was acidified with conc. HCl to pH 5 and the solids formed were filtered off. The solids were dried overnight and purified via column chromatography (0 to 20% MeOH/EtOAc) to afford the desired product in 74% yield (2.23 g). ^1^H NMR (400 MHz, DMSO-*d*6): δ ppm 1.19 (t, *J* = 7.6 Hz, 3H), 2.69 (q, *J* = 7.6 Hz, 2H), 3.68 (s, 3H), 10.07 (s, 1H), 11.84 (s, 1H). LC-MS m/z (M + H)^+^ = 195.

#### 4,6-dichloro-3-ethyl-1-methyl-1*H*-pyrazolo[3,4-*d*]pyrimidine (IV)

To a round bottom flask containing 3-ethyl-1-methyl-1*H*-pyrazolo[3,4-*d*]pyrimidine-4,6(5*H*,7*H*)-dione (I**II**, 500 mg, 2.57 mmol) was added phosphorus (V) chloride (1 g, 5.14 mmol) and phosphoryl chloride (10 mL). The mixture was refluxed for 9 h and then allowed to cool room temperature. After POCl3 was removed by rotary evaporation, the solid was washed with water, and the yellow solid residue was purified by flash chromatography on silica gel (EtOAc/Hexane) to provide pure white solid intermediate compound (185 mg, 31%). LC-MS *m/z* 231 (MH)^+^. ^1^H NMR (400 MHz, DMSO-*d*6): δ ppm 1.34 (t, *J* = 7.2 Hz, 3H), 3.06 (q, *J* = 7.2 Hz, 2H), 3.96 (s, 3H).

#### 6-chloro-3-ethyl-N-(4-fluorobenzyl)-1-methyl-1H-pyrazolo[3,4-d]pyrimidin-4-amine

(V). 4,6-dichloro-3-ethyl-1-methyl-1H-pyrazolo[3,4-d]pyrimidine (**IV**, 1.0 g, 4.33 mmol) was dissolved in 10 mL acetonitrile before adding DIPEA (1.10 mL, 6.50 mmol) and 4-fluorobenzyl amine (0.80 mL, 6.50 mmol). The resulting solution was stirred at room temperature for 6 hrs and concentrated. The crude product was purified via silica gel chromatography (20% MeOH/EtOAc) to afford the desired product in 65% yield (897 mg). ^1^H NMR (400 MHz, DMSO-*d*6) δ 8.09 (t, *J* = 5.87 Hz, 1H), 7.40 (dd, *J* = 5.62, 8.56 Hz, 2H), 7.15 (t, *J* = 8.80 Hz, 2H), 4.69 (d, *J* = 5.99 Hz, 2H), 3.78 (s, 3H), 2.98 (q, *J* = 7.46 Hz, 2H), 1.23 (t, *J* = 7.46 Hz, 3H). LC-MS m/z (M + H)^+^ = 320.

#### Ethyl 4-(3-ethyl-4-((4-fluorobenzyl)amino)-1-methyl-1H-pyrazolo[3,4-d]pyrimidin-6-yl)benzoate

6-chloro-3-ethyl-N-(4-fluorobenzyl)-1-methyl-1H-pyrazolo[3,4-d]pyrimidin-4- amine (**V**, 300 mg, 0.94 mmol), (4-(ethoxycarbonyl)phenyl)boronic acid (327.8 mg, 1.69 mmol), Pd(dppf)Cl2 (95.1 mg, 0.13 mmol), K2CO3 (298.5 mg, 2.16 mmol), 12 mL DMF and 3 mL H2O were weighed into a 20 mL microwave vial and capped. The vial was vacuumed and backfilled with Argon gas. This process was repeated 3X and the vial kept under positive Argon pressure. Anhydrous DMF and degassed distilled water were added to the vial in a 4:1 ratio. The microwave vial was placed in a heating block and the block heated to 100 °C overnight with stirring. After this time, the reaction was cooled to room temperature and filtered through a short pad of Celite eluting with ethyl acetate. The organic was transferred into a separatory funnel containing water. The aqueous layer was extracted two times with ethyl acetate. All the organics were combined, washed with brine, dried over magnesium sulfate, and filtered. The filtrate was concentrated, and the crude product purified via silica gel chromatography eluting with hexanes and ethyl acetate. To ensure sufficiently pure compounds to submit for bioassay, a second reversed phase chromatography eluting with water and acetonitrile with no modifiers was performed. The product was isolated in 56% yield as an off-white solid (226 mg). ^1^H NMR (400 MHz, DMSO-*d*6) δ 8.49 (d, *J* = 8.44 Hz, 2H), 8.05 (d, *J* = 8.44 Hz, 2H), 7.89 (s, 1H), 7.51 (dd, *J* = 5.69, 8.50 Hz, 2H), 7.15 (t, *J* = 8.93 Hz, 2H), 4.85 (d, *J* = 5.50 Hz, 2H), 4.35 (q, *J* = 7.09 Hz, 2H), 3.92 (s, 3H), 3.05 (q, *J* = 7.46 Hz, 2H), 1.35 (t, *J* = 7.09 Hz, 3H), 1.29 (t, *J* = 7.46 Hz, 3H). LC-MS m/z (M + H)^+^ = 434.

#### 4-(3-ethyl-4-((4-fluorobenzyl)amino)-1-methyl-1*H*-pyrazolo[3,4-*d*]pyrimidin-6- yl)benzoic acid (PDEi5)

Ethyl 4-(3-ethyl-4-((4-fluorobenzyl)amino)-1-methyl-1*H*- pyrazolo[3,4-*d*]pyrimidin-6-yl)benzoate (100 mg, 0.231 mmol) was weighed into a round bottom flask before adding KOH (23.3 mg, 0.347 mmol) and 10 mL ethanol. The flask was placed in a heating block and heated to 100 °C for 2 h with a condenser on top. After this time, the reaction was cooled to room temperature, acidified with 1 M HCl solution to pH 6 and concentrated. The resulting crude product was purified via reversed phase chromatography eluting with H2O:CH3CN. The product was isolated in 73% yield as a white solid (68 mg). ^1^H NMR (400 MHz, DMSO-*d*6) δ 13.03 (br. s., 1H), 8.48 (d, *J* = 8.31 Hz, 2H), 8.03 (d, *J* = 8.31 Hz, 2H), 7.92 (t, *J* = 5.87 Hz, 1H), 7.51 (dd, *J* = 5.81, 8.25 Hz, 2H), 7.15 (t, *J* = 8.80 Hz, 2H), 4.86 (d, *J* = 5.50 Hz, 2H), 3.91 (s, 3H), 3.05 (q, *J* = 7.46 Hz, 2H), 1.28 (t, *J* = 7.46 Hz, 3H). LC-MS m/z (M + H)^+^ = 406. HRMS (ESI) m/z: [M + H]^+^ Calcd for C22H20FN5O2 406.1679; found 406.1670.

### Cell culture and parasites

Human ileocecal adenocarcinoma (HCT-8) cells (ATCC) were cultured in RPMI 1640 medium (Invitrogen) supplemented with 10% heat-inactivated fetal bovine serum (Sigma-Aldrich), 120 U/ml penicillin, and 120 μg/ml streptomycin (ATCC) at 37°C and 5% CO2. HCT- 8 cells were used between passages 9 and 39 for all experiments. *C. parvum* Iowa strain oocysts were purchased from Bunch Grass Farm (Deary, ID), and sorted in phosphate-buffered saline (PBS) with penicillin and streptomycin at 4°C for up to 5 months prior to use. *C. hominis* TU502 strain oocysts were purchased from the Tzipori laboratory (Tufts University) and used within 1 month of acquisition.

### Immunofluorescence assay for drug screening and follow-up growth inhibition assays

The same assay method was used to measure both *C. parvum* and *C. hominis* growth, except for adjustment in the inoculum size used to account for reduced *C. hominis* infectivity. The method was as described previously^27^. Oocysts were excysted by treatment with 10 mM hydrochloric acid (10 min, 37°C), followed by exposure to 2 mM sodium taurocholate (Sigma-Aldrich) in PBS (10 min, 16°C). Excysted oocysts were then added to >95% confluent HCT-8 cell monolayers in 384-well plates. *C. parvum* and *C. hominis* oocysts were used at 5,500 per well. Drug compounds were added 3 h post infection for both screening and follow-up assays and parasites were allowed to grow for 48 h post infection, at which time the plates were washed three times with PBS, fixed with 4% paraformaldehyde in PBS (15 min, room temperature), permeabilized with 0.25% Triton X-100 (10 min, 37°C), washed three times with PBS with 0.1% Tween 20, and blocked with 4% bovine serum albumin (BSA) in PBS for 2 h at 37°C or overnight at 4°C. Parasitophorous vacuoles were then stained ^63^ with 1.33 μg/ml fluorescein-labeled *Vicia villosa* lectin (Vector Laboratories) diluted in 1% BSA in PBS with 0.1% Tween 20 (1 h, 37°C), followed by addition of Hoechst 33258 (AnaSpec) at a final concentration of 90 μM in water (15 min, 37°C). Wells were then washed five times with PBS with 0.1% Tween 20. Images were acquired using a Nikon Eclipse TE2000 epifluorescence microscope (Nikon, USA) with an automated stage programmed to acquire a 3 × 3 tiled image of the center of each well using an Exi Blue fluorescence microscopy camera (QImaging, Canada) and a 20x objective (numerical aperture 0.45). The nucleus and parasite image channels were analyzed using NIH ImageJ (National Institutes of Health) and previously published macros^27^.

### *Cryptosporidium* time-kill assays

Excysted *C. parvum* oocysts were added to >90% confluent HCT-8 cells in 384-well plates. Compounds were added after 24 h of infection at concentrations in multiples of the EC90. At 24 h after infection (i.e. time of compound addition) and the shown time-intervals thereafter, the plates were washed, fixed, and prepared for parasite enumeration using epifluorescence microscopy as described for the growth inhibition assay described above. A separate 384-well plate was used for each time point. Parasite numbers were normalized to host cell nucleus numbers and dimethyl sulfoxide (DMSO) controls on each assay plate to determine the effect of compound inhibition at each time point.

### Assays to determine the life-stage affected by drug compounds

Assays to assess the life-cycle stage affected by inhibitors were all performed as in Jumani, et al. and using compounds at 2x the EC90 measured in the *C. parvum* growth assay^46^. Each of the assays used *C. parvum* oocysts induced to excyst and infection of HCT-8 cell monolayers as described above (see assay for drug screening and growth inhibition). Modifications made to the protocol to assay each life-cycle stage were as follows.

To assay host cell invasion, HCT-8 cell monolayers were pre-treated with 2 x the EC90 of each compound and incubated for 1 h prior to infection. Oocysts induced to excyst with HCl and Na^+^ taurocholate were added and invasion was allowed to proceed for 3 hours at 37°C before washing with PBS with 111 mM D-galactose to remove extracellular parasites. The monolayers were then fixed, permeabilized, and stained for microscopy as above to enumerate the parasitophorous vacuoles.

Because *C. parvum* infection in the HCT-8 system begins crudely syncrhonized, the number of parasitophorous vacuoles ratchets up with each round of asexual replication, cell egress, and invasion of new cells^46^. Therefore, to quantify *C. parvum* egress and re-invasion of host cells, HCT-8 monolayers grown in 384-well plates were infected and exposed to compounds as for the growth assay, and the number of parasite vacuoles was measured at 6 h and 19.5 h following infection, which are time points bracketing the time required for the first round of asexual replication. Data are then expressed as the ratio of parasite vacuoles at 19.5 h and 6 h and viewed relative to the vehicle control and the known egress inhibitor MMV403679^46^.

Finally, development of female gametocytes was assayed by measuring the percent of *C. parvum* that stain for the female gametocyte marker DNA meiotic recombination protein (DMC1) which typically is expressed in female gametocytes at ∼ 48 h in the HCT-8 culture system^46^. HCT- 8 cells were infected and treated as for the growth assay, except that compounds were added at 30 h post-infection to exclude early effects on asexual replication. The percent of parasites expressing DMC1 at 48 h was then determined by staining all parasite vacuoles with *V. villosa* lectin and staining DMC1 with an anti-*C. parvum* DMC1 mouse monoclonal antibody (clone 1H10G7 (IgG2b, kappa) used as undiluted culture supernatant at 25 μl/well) and a secondary Alexa Fluor 568 goat anti-mouse IgG antibody (Invitrogen, catalog# A-11004) at 1:500 dilution (4 µg per mL).

### Host cell cytotoxicity assays

Host cell toxicity was measured using CellTiter AQueous assay (Promega, USA) according to the manufacturer’s instructions. This method measures optical absorbance to measure reduction of the tetrazolium compound [3-(4,5-dimethylthiazol-2-yl)-5-(3-carboxymethoxyphenyl)-2-(4-sulfophenyl)-2H-tetrazolium, inner salt; MTS] by NADH and NADPH produced by dehydrogenases in viable cells. Assays were conducted using >95% confluent HCT-8 cells grown in 384-well plates for 48 h (37°C) with varying concentrations of compounds. The corner wells of each plate were trypsinized to remove cells and were used as blanks for measuring absorbance at 490 nm, and data were expressed as the percentage of cell viability for vehicle controls on each assay plate (DMSO). GraphPad PRISM software, version 9.4.1, was used to calculate the 50% toxic concentration (TC50) and selectivity indices were determined as the ratio of *C. parvum* EC50 to TC50.

### NOD SCID gamma mouse model of cryptosporidiosis

All mouse experiments were performed in compliance with animal care guidelines and were approved by the University of Vermont Institutional Animal Care and Use Committee. NOD SCID gamma mice (NOD.Cg-*Prkdc^scid^ Il2rg^tm1Wjl^*/SzJ) were purchased from The Jackson Laboratory (Bar Harbor, ME, USA) and housed for at least a week for acclimatization. At the age of 4 to 5 weeks, mice were infected with 10^5^ *C. parvum* oocysts and compounds were dosed by oral gavage beginning 7 days post-infection according to the regimen specified for each experiment (four mice per experimental group). Aliquoted DMSO stock solutions were stored for < 10 days at -80°C. Compound doses were prepared on the day of dosing by thawing and sonicating an aliquot of the DMSO stock, followed by dilution in 1% hydroxypropyl methylcellulose (HPMC) to a final concentration of 5% DMSO in 100 μl of 1% HPMC per dose.

Oocyst shedding in feces was monitored at the specified time points qPCR using a previously described protocol^64^.

### Pharmacokinetic studies

Both intravenous and oral pharmacokinetic studies were conducted using 6–8-week-old male CD-1 mice (Charles River Labs). Prior to each study, they were housed in a temperature- and light-controlled environment for 5 days. All mice had free access to standard feed and water prior to and throughout the study. All procedures were performed in accordance with the Institutional Animal Care and Use Committee of Saint Louis University. Intravenous dosing solution was prepared the day of the study in 60% Saline/40% Formal Glycerol to a final concentration of 0.24 mg/mL, which resulted in a clear solution. Drug was administered into the tail vein (3 mice/timepoint) at 1 mg/kg. At the given time point (0.08, 0.17, 0.5, 1, 2, 4, 6 hr), the mouse was euthanized by CO2 exposure and samples collected. Oral administration of drug (2.4 mg/mL, 10 mg/kg) was performed with a solution of 0.5% Tween 80/0.5% Carboxymethyl cellulose in water. Drug was given by mouth using a stainless-steel feeding needle (3 in x 2.9 mm ball). At the given time point (0.25, 0.5, 1, 2, 4, 7, 24 hr), the mouse was euthanized by CO2 exposure and samples collected. In both studies, blood was obtained by cardiac puncture into sodium heparin tube (vacutainer) and placed on ice. The blood was separated by centrifugation and the plasma stored at -80°C until analysis. Plasma proteins were denatured by the addition of 3 volumes of ice-cold acetonitrile containing internal standard (5 ng/ml Enalapril). Plasma samples were then vortexed and centrifuged at 12,000xg for 10 minutes. Supernatants were then transferred to a 96-well sample plates for analysis.

The intestinal tissue was obtained by dissection. Briefly, after exposing the abdominal cavity, the entire length of the intestine was removed and 4 cm segments excised from the beginning (duodenum), middle (jejunum), and end (ileum) of the small intestine, and from the proximal colon. The segments were sliced along the longitudinal axis, and the contents thoroughly rinsed away with sterile water. The tissues were stored at -80°C until futher processed for analysis.

Intestinal tissue was macerated in a bead beater in PBS, pH 7.2. Tissue mascerates were centrifuged at 12,000g for 10 minutes. The supernate was then treated with 3 volumes of ice-cold acetonitrile containing internal standard (5 ng/ml Enalapril) followed by centrifugation.

The final supernate was then transferred to a 96-well sample plate for analysis.

Both plasma and tissue samples were analyzed by liquid chromatography-tandem mass spectrometry (LC/MS/MS), using compound spiked into control plasma as a standard.

Noncompartmental analysis was performed using Phoenix WinNonlin version 8.2 (Certara, Clayton, MO).

### *In silico* sequence analysis and docking studies

Putative *C. parvum* PDEs were identified using CryptoDB and BLAST to search for genes encoding proteins homologous to human PDE5A (NCBI sp|076074)^65,66^. CryptoDB genes cgd3_2320 and cgd6_4020 were designated as *Cp*PDE1 and *Cp*PDE2, respectively. AlphaFold *Cp*PDE models (AlphaFold: A3FQ29-F1-model_v4 (*Cp*PDE1) and A3FPW6-F1-model_v4 (*Cp*PDE2)) were accessed on August 22, 2023 (last updated in AlphaFold DB version 2022-11- 01)^41,42^. The crystal structure of a *Schistosoma manosoni* PDE (*Sm*PDE4 (pdb 6EZU^44^)) was overlayed to enable addition of active site metal ions. Merged AlphaFold/metal ion structures were minimized using the Protein Preparation panel in Schrödinger Maestro to converge heavy atoms to RMSD 0.30 Å using the OPLS4 forcefield. Schrödinger Maestro and the SP Glide docking routine were used to produce structures for molecular docking studies^45^. The models were further refined using the Induced Fit docking routine in Schrödinger Maestro with and without active site waters.

### SafetyScan47 assay panel, mammalian phosphodiesterase assay panel, CYP inhibition panel, hERG binding and aqueous solubility

Potential off target effects of PDEi5 were screened in 78 assays (47 targets) with the Eurofins SafetyScan47 panel^47^. Assay technologies used include: cAMP secondary messenger assays using cell lines expressing non-tagged GPCRs, a calcium secondary messenger assay to measure GPCR activity in live cells, a nuclear hormone receptor assay (NHR Pro) and Nuclear Translocation (NHR NT) assays, a proprietary kinase screening platform (KINOMEScan) that uses competition to quantify kinase binding, a neurotransmitter uptake assay, potassium and membrane potential assays, and specific enzymatic assays used to measure consumption of a substrate or production of a product. Mammalian PDE inhibition assays were also performed by Eurofins, using PDEs sourced as indicated in Table S5. The assay method for each enzyme was to measure production of tritiated product (either [^3^H]-5’GMP or [^3^H]-5’AMP (substrates indicated in Table S5)) by scintillation counting. Results were expressed as a percent inhibition of control specific activity in the presence of 10 μM PDEi1. CYP inhibition was measured by Eurofins using the indicated recombinant human enzymes. hERG affinity was measured by Eurofins at 10 µM as % inhibition of the binding of [^3^H]-dofetilide to the human potassium channel hERG. Aqueous solubility was measured by Eurofins as kinetic solubility in aqueous solution at pH 7.4.

### Statistical information, and figure preparation

GraphPad Prism software (version 9.5.1) was used to prepare all graphs, calculate compound dose-response curves, calculate decay curves for time-kill assays, and perform statistical analyses. The limit of detection of the qPCR assay performed on mouse feces is ∼ 100 oocysts per gram of dried feces, and samples for which no signal was detercted were plotted at the LOD for purposes of graphing and statistical analysis. Statistical significance was assessed as indicated in figure legends. Graphs were exported as .eps files, and final figures were prepared using Adobe Illustrator.

## Acknowledgments

We thank Merck KGaA, Darmstadt, Germany and the Open Innovation Portal for generous provision of compounds for screening and follow up studies, and for generous sharing of pre-existing data. CDH, MJM, and DWG were funded by NIAID R21R33 AI141184.

---

**Supplemental Figure 1:**
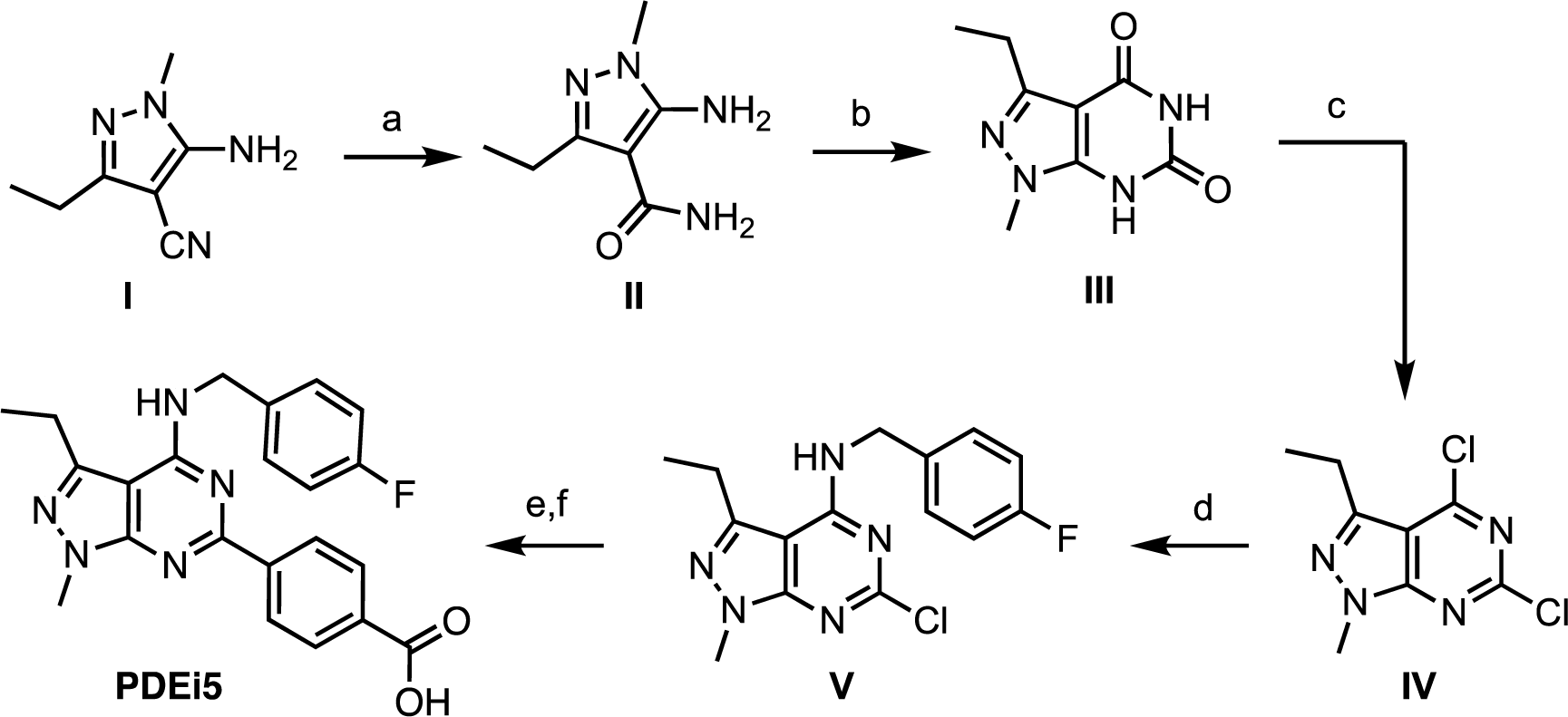
Synthesis of PDEi5. Reassgents and conditions: (a) conc. H2SO4, 25 °C, overnight, 56%; (b) urea, 200 °C, 2 h, 74%; (c) PCl5, POCl3, 110 °C, overnight, 38%; (d) 4- fluorobenzylamine, DIPEA, acetonitrile, 25 °C, 65%; (e) (4-(ethoxycarbonyl)phenyl)boronic acid, Pd(dppf)Cl2, K2CO3, DMF:H2O (4:1), 110 °C, overnight, 56%; (f) KOH, Ethanol, 100 °C, 2 h, 73%.

**Supplemental Figure 2:**
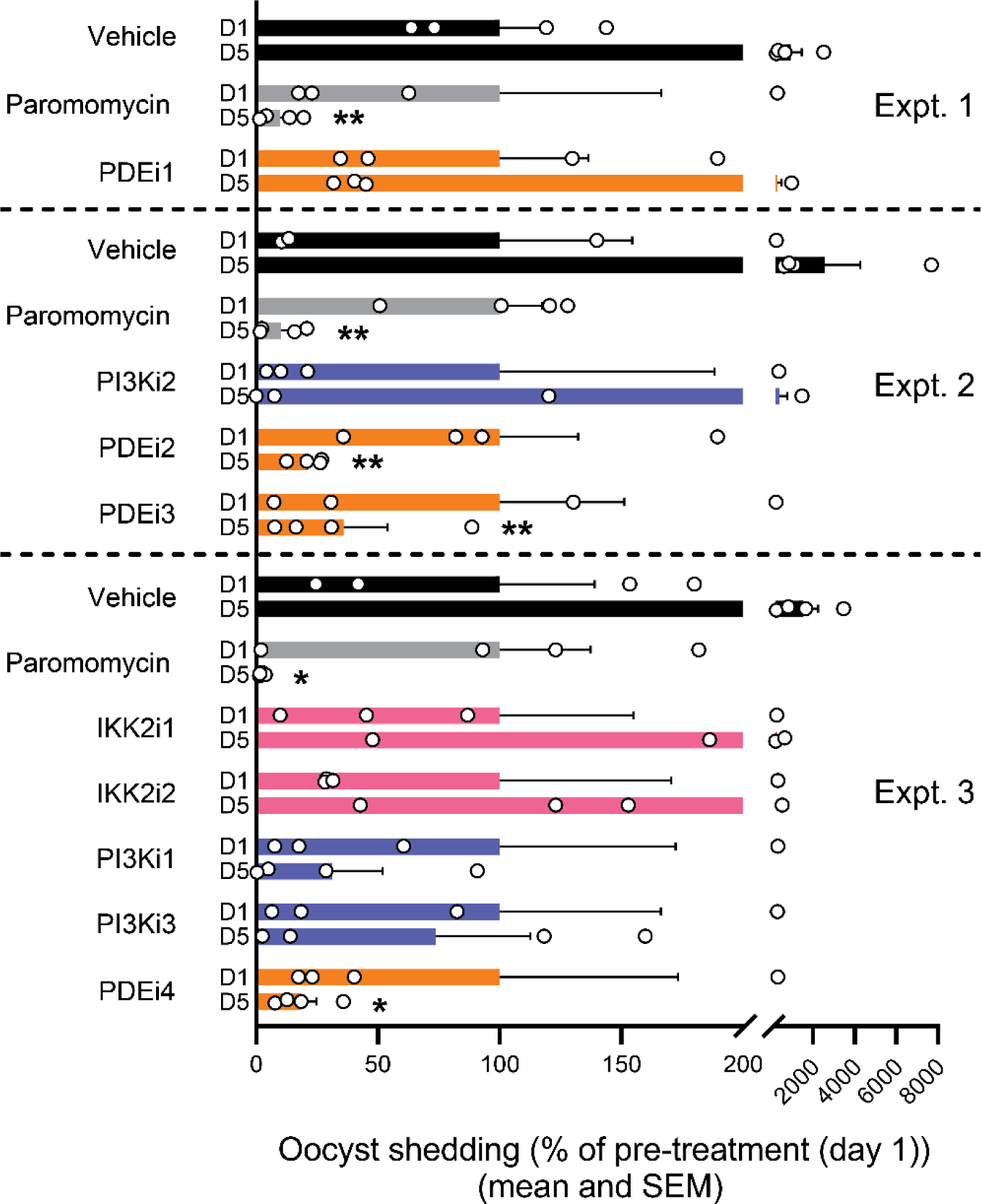
NSG mouse experiments for preliminary drug lead identification. The indicated compounds were tested for efficacy in established cryptosporidiosis in NSG mice. *C. parvum* infection was allowed to progress for 7 days prior to administering 50 mpk twice daily of the indicated compounds or 1000 mpk twice daily of paromomycin (positive control) by oral gavage for 4 days. D1 or D5 indicates day after first drug dose. Parasite shedding per mg of feces was determined by qPCR. Data are shown as the percent of pre-treatment parasite shedding for each experimental group (mean and SEM; n = 4 mice per experimental group). Asterisk (*) indicates p = 0.05 and double asterisk (**) p = 0.14 vs. vehicle control for the indicated experiment by one-way ANOVA with Dunnett’s multiple comparisons test.

**Supplemental Figure 3:**
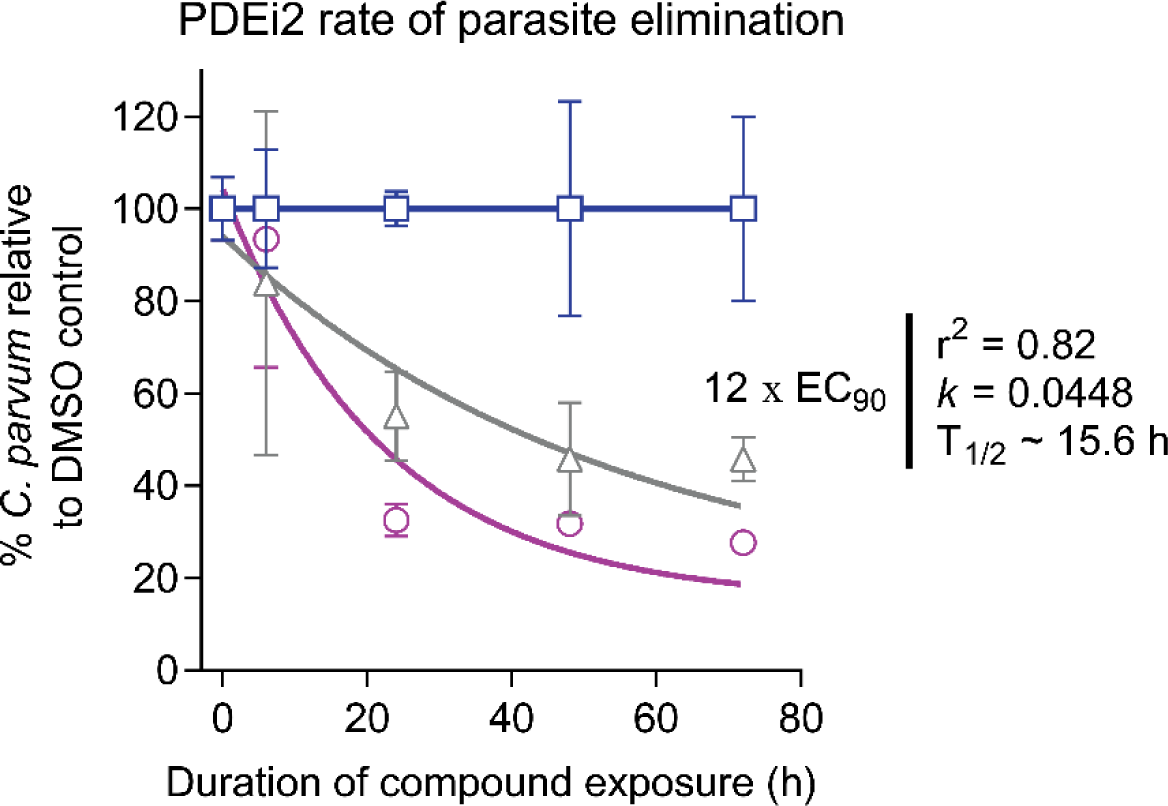
Time-kill curve for PDEi2 showing elimination of *C. parvum* from infected HCT-8 cells. One-phase exponential decay curves fit for the highest concentration of PDEi2 tested. Data are the means and SD for 4 culture wells per time point, normalized for each time point to the vehicle control (DMSO). Curves are shown for 3×EC90 (grey), 12×EC90 (cyan), or the DMSO control (blue).

**Supplemental Figure 4:**
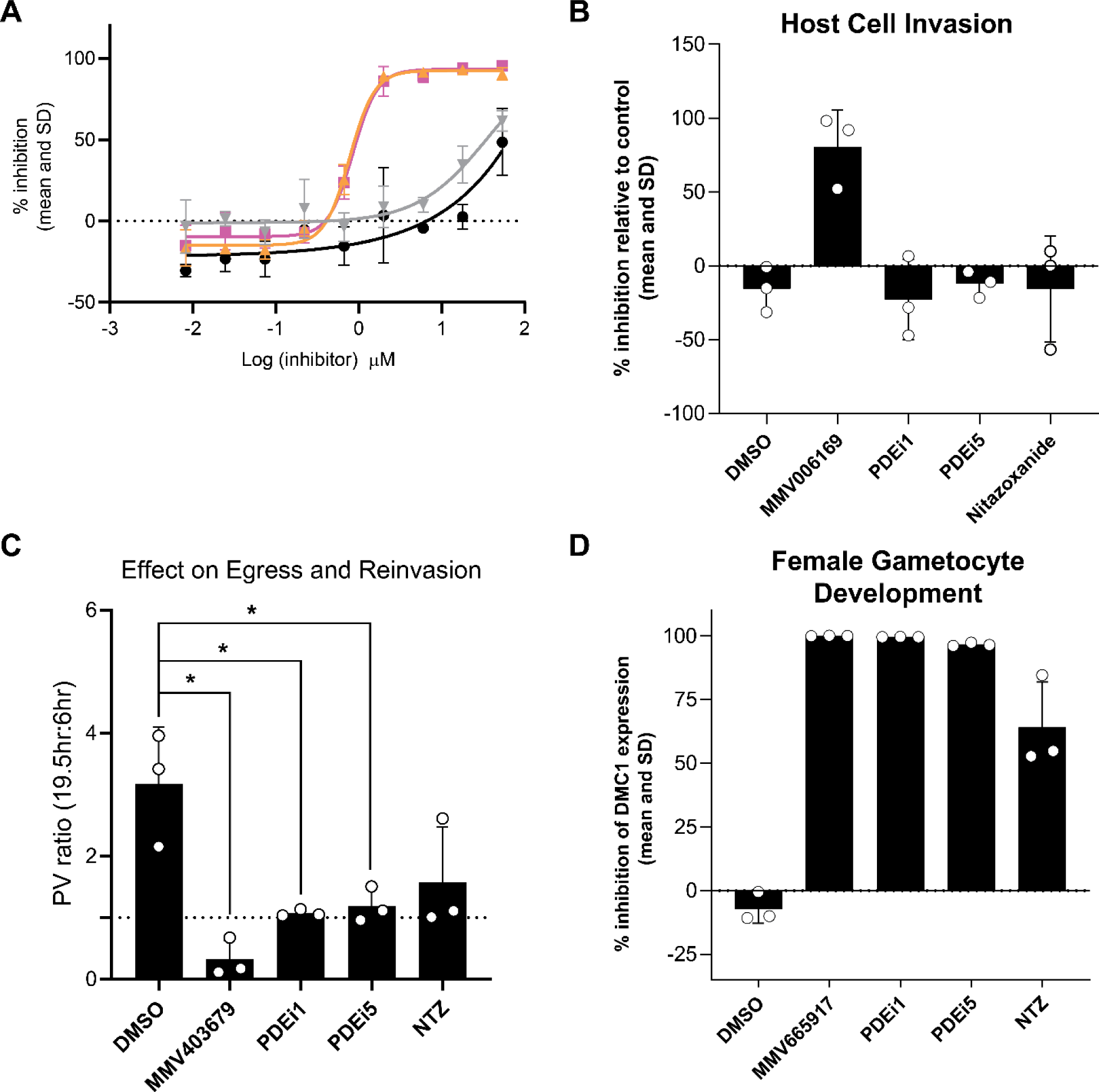
Phenotypic basis of the anticryptosporidial effect of new PDEis. (A) Effect of new PDEi series on *C. parvum* growth compared to other potent mammalian PDE5 inhibitors. Dose response assays were conducted against *C. parvum* (Iowa) grown in HCT-8 cells. Data are mean and SD for two biological replicates. Cyan = PDEi2; orange = PDEi1; grey = Cmp7a; black = sildenafil. (B) PDE inhibitors have no effect on host cell invasion by *C. parvum.* Compounds were added prior to addition of excysted parasites, and intracellular parasites were assayed after 3 h. The known invasion inhibitor MMV006169 was used as a positive control. (C) Effect of the PDEi series on *C. parvum* host cell egress and reinvasion. Cell egress and reinvasion were assessed by determining the ratio of parasite vacuoles before and after the first cycle of asexual replication, and results were compared to vehicle or the known egress inhibitor MMV403679. (D) PDE inhibitors block *in vitro* sexual development of *C. parvum.* Compounds were added 30 hours after infection of HCT-8 cell monolayers, and expression of the macrogametocyte marker DMC1 was measured at 48 h. The percentage of DMC1-expressing parasites following treatment with the known inhibitor MMV665917 was used as a positive control. (B), (C), and (D) All compounds were assayed at 2× the EC90 for growth inhibition. All data are the means from three individual biological replicates. * indicates p < 0.01 for one-way ANOVA with Dunnett’s multiple comparisons test.

**Supplemental Figure 5:**
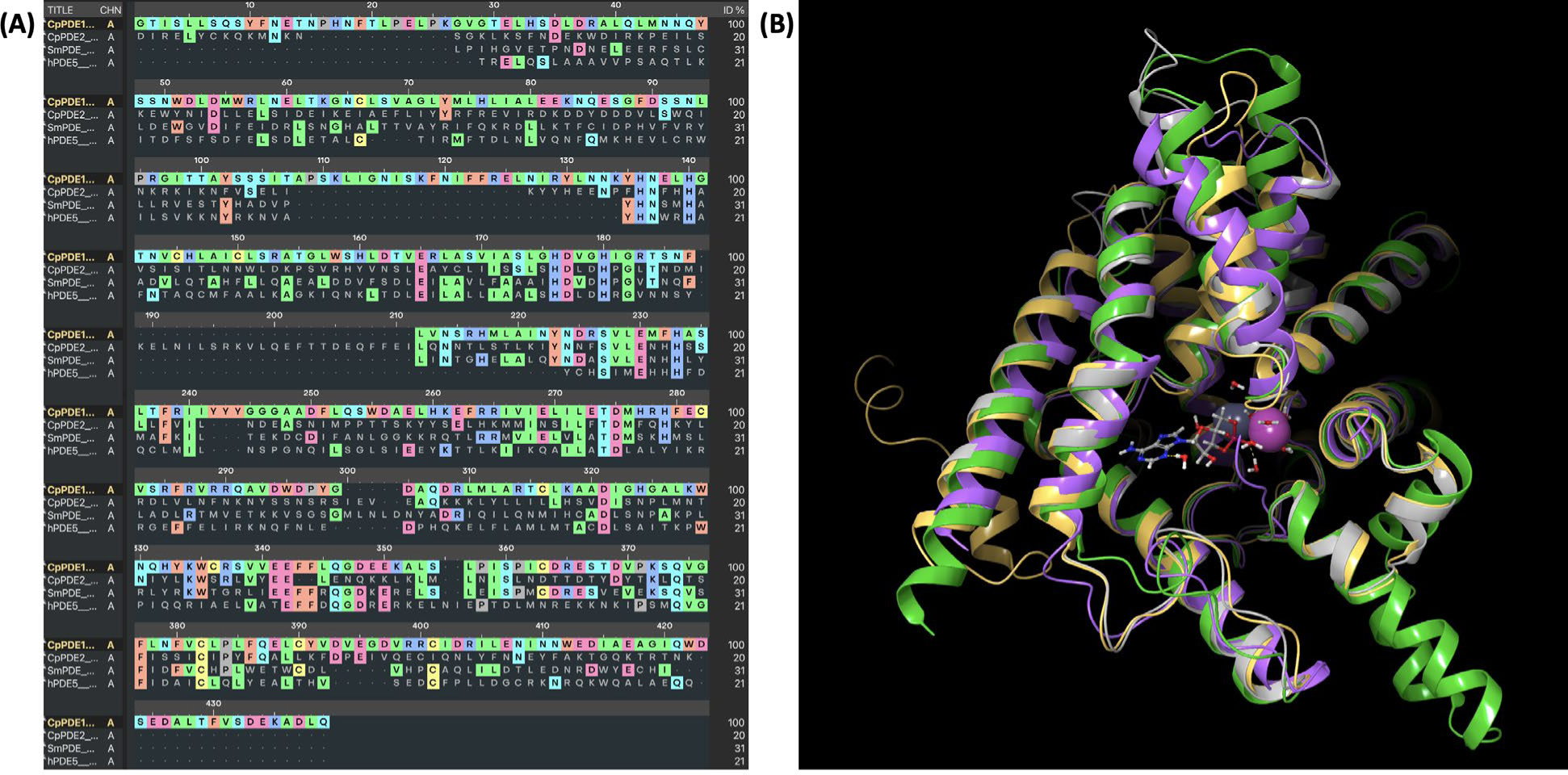
(A) Sequence alignment of catalytic domains of CpPDE1, CpPDE2, SmPDE, and hPDE-V. (B) Overlay of hPDE-V (pdb 1UDT; purple), SmPDE with bound cAMP (pdb 6EZU; gray), CpPDE1 model (gold), and CpPDE2 model (green).

**Supplemental Figure 6:**
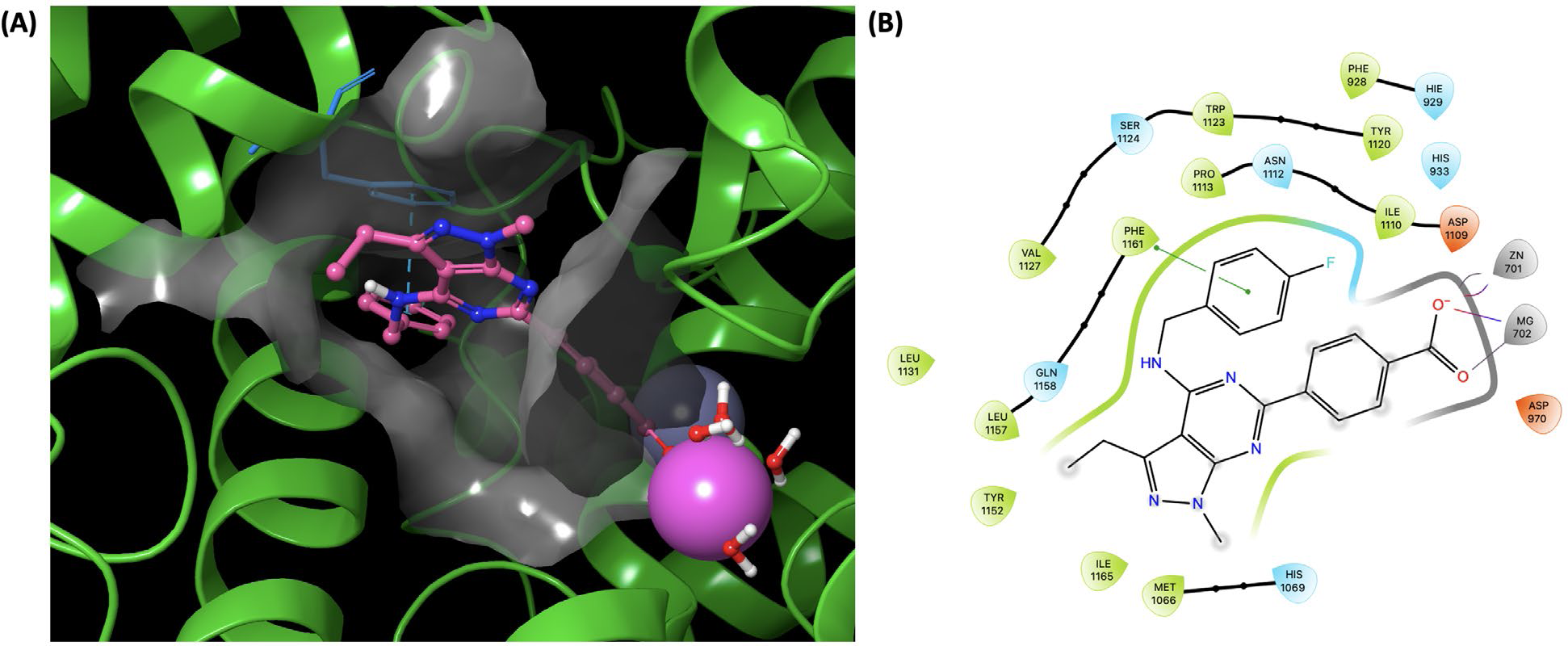
Lowest energy pose from induced fit docking of PDEi5 in the CpPDE2 model. Glide docking score -11.7. The ligand binding pocket of CpPDE2 is shown as a gray surface. (A) Binding site illustrating Phe935 (blue) forming π-π face-face interactions with PDEi5. (B) Ligand interaction diagram illustrating residues within 4Å of the ligand, π-stacking between ligand and Phe935 (green line), and ionic salt bridge and hydrogen bonding between the carboxylate of the ligand with the zinc and magnesium.

**Supplemental Figure 7:**
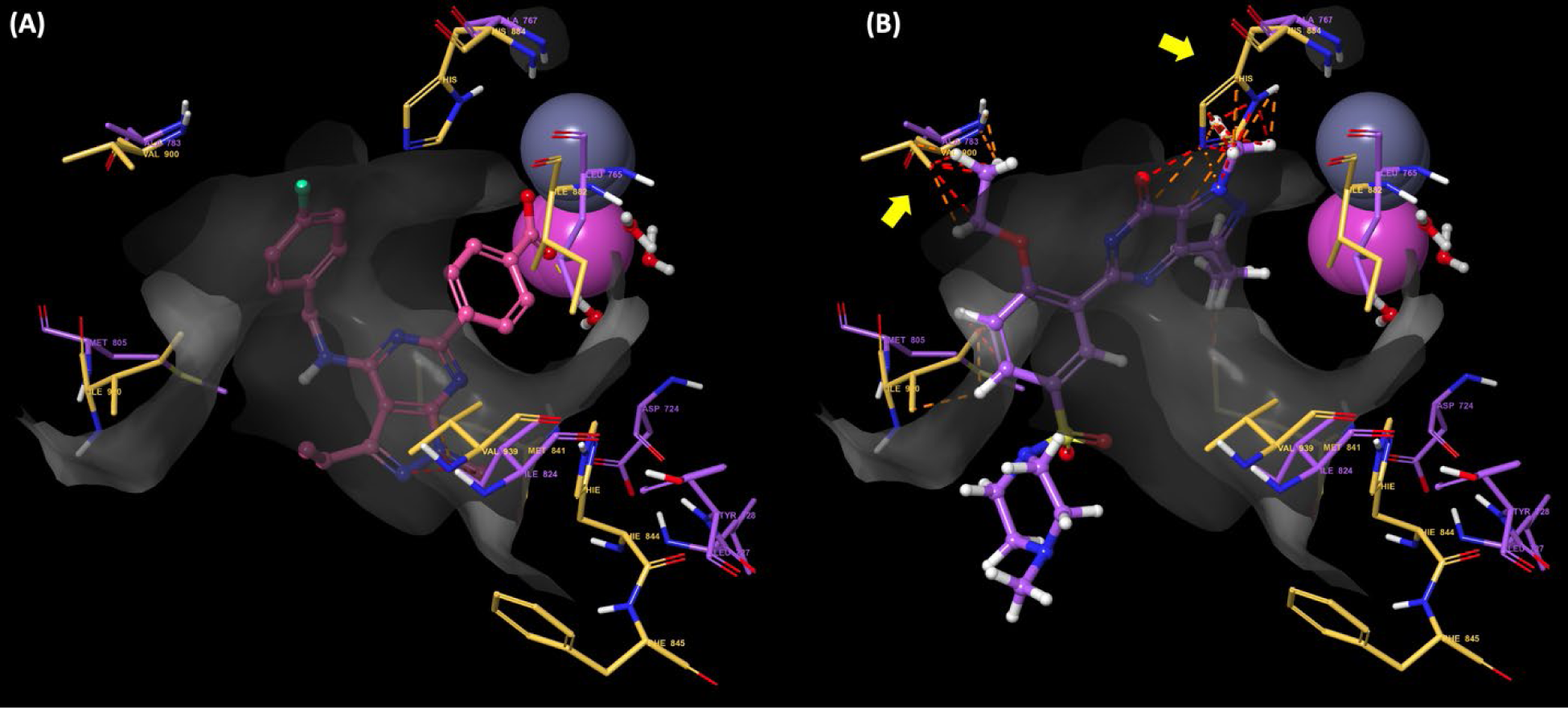
Overlay of lowest energy pose from induced fit docking of PDEi5 in the *Cp*PDE1 model (gold) with hPDE-V (purple; pdb 1UDT). Only the seven of the 18 total residues within 4Å of the PDEi5 ligand that differ between the two proteins are shown, along with Val900 (5Å). The ligand binding pocket of *Cp*PDE1 is shown as a gray surface. (A) Induced fit docked pose of PDEi5. (B) Overlay of sildenafil from hPDE-V structure (pdb 1UDT). Yellow arrows indicate steric clashing preventing sildenafil from obtaining the same docking pose in *Cp*PDE1 vs. hPDE-V.

**Supplemental Figure 8:**
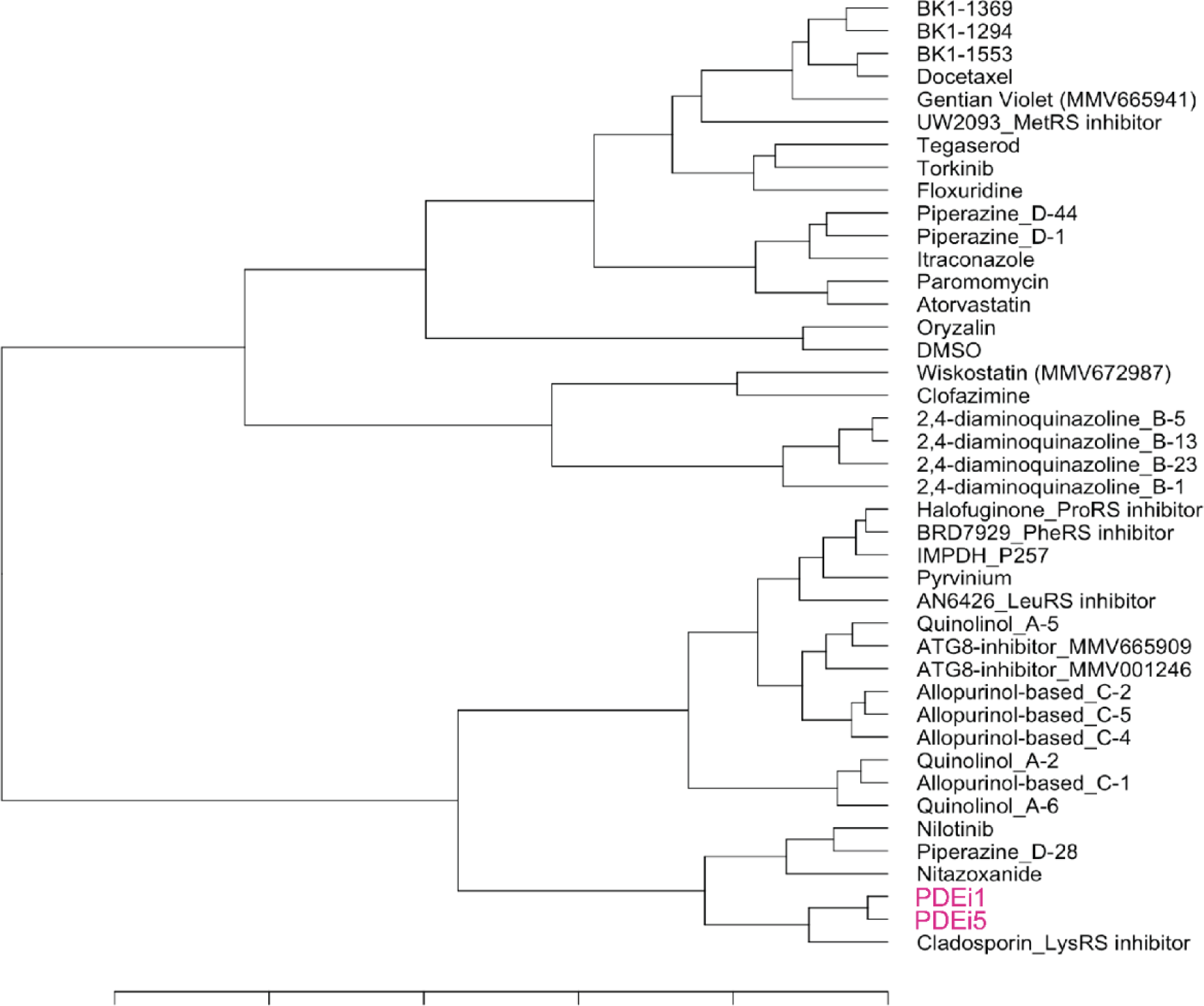
Dendrogram generated from phenotypic assay data for phosphodiesterase inhibitors and publicly available data for anticryptosporidial lead compounds.

**Supplemental Figure 9:**
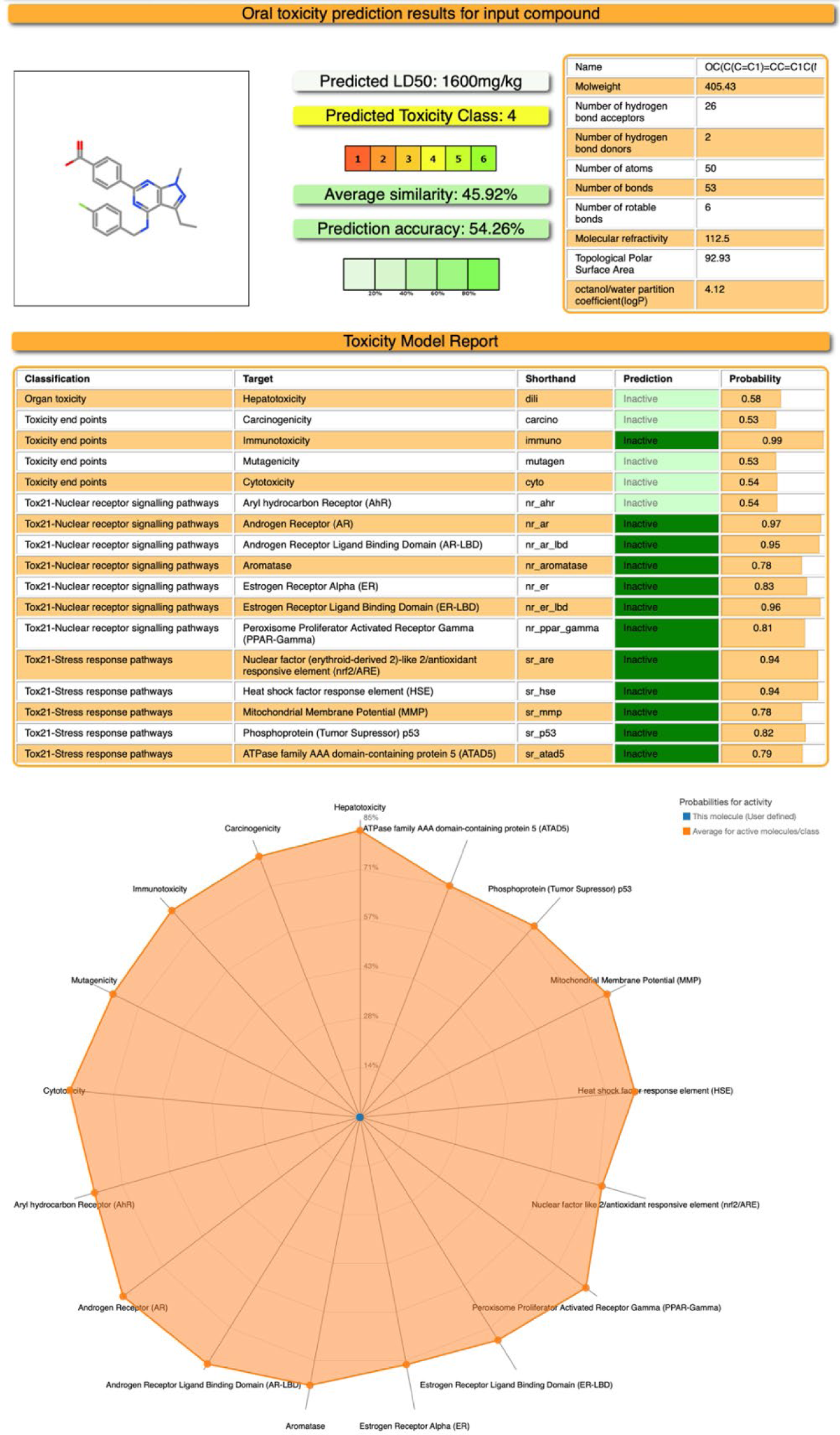
In silico toxicity prediction for PDEi5. Data generated by ProTox-II (https://tox-new.charite.de/protox_II/). Direct access at https://tox-new.charite.de/protox_II/index.php?site=direct_load&id=16927400934198&models=dili,carcino,immuno,mutagen,cyto,nr_ahr,nr_ar,nr_ar_lbd,nr_aromatase,nr_er,nr_er_lbd,nr_ppar_gamma,sr_are,sr_hse,sr_mmp,sr_p53,sr_atad5

**Table S1:** Characteristics of compounds tested *in vivo*. *In vitro and in vivo* PK/ADME data. Green coding indicates typically desired property, yellow indicates moderate, and red indicates typically undesired property.

| Chemotype | Cmpd ID | Cp EC<sub>50</sub> (μM) | Ch EC<sub>50</sub> (μM) | HCT8 TC<sub>50</sub> (μM) | MLM CI (μg/mL/min) | Mouse t<sub>1/2</sub> (h) | %F | Log P | Caco Efflux |
| --- | --- | --- | --- | --- | --- | --- | --- | --- | --- |
| Aminopyrimidine | IKK2i1 | 5.4 | n.d. | n.d. | 27 | 2-3 | n.d. | 4.11 | 1.5 |
| Pyrazole | IKK2i2 | 0.020 | n.d. | n.d. | 572 | 0.25 | <5 | 3.16 | 158 |
| Thiazolidinedione | PI3Ki1 | 0.14 | n.d. | n.d. | 22 | 3 | 100 | 1.77 | 1.5-1.9 |
|  | PI3Ki2 | 0.13 | n.d. | n.d. | 77 | n.d. | n.d. | 2.51 | 1.6 |
|  | PI3Ki3 | 0.16 | n.d. | n.d. | 138 | 1.2 | ~46 | 2.33 | n.d. |
| Benzothieno-pyrimidine | PDEi1 | 0.98 | n.d. | n.d. | n.d. | 0.3 | <5 | 6.66 | n.d. |
| Pyrazolopyrimidine | PDEi2 | 0.86 | 0.61 | >100 | <10 | 0.9 | 33 | 5.29 | n.d. |
| Benzothieno-pyrimidine | PDEi3 | 1.0 | n.d. | n.d. | n.d. | 0.7 | ~15 | 7.47 | 0.6-1.6 |
| Indolopyrimidine | PDEi4 | 0.87 | 1.9 | 87 | n.d. | n.d. | n.d. | 6.55 | n.d. |
| Pyrazolopyrimidine | PDEi5 | 1.8 | 4.7 | >100 | n.d. | n.d. | n.d. | 4.7 | n.d. |

**Table S2:**
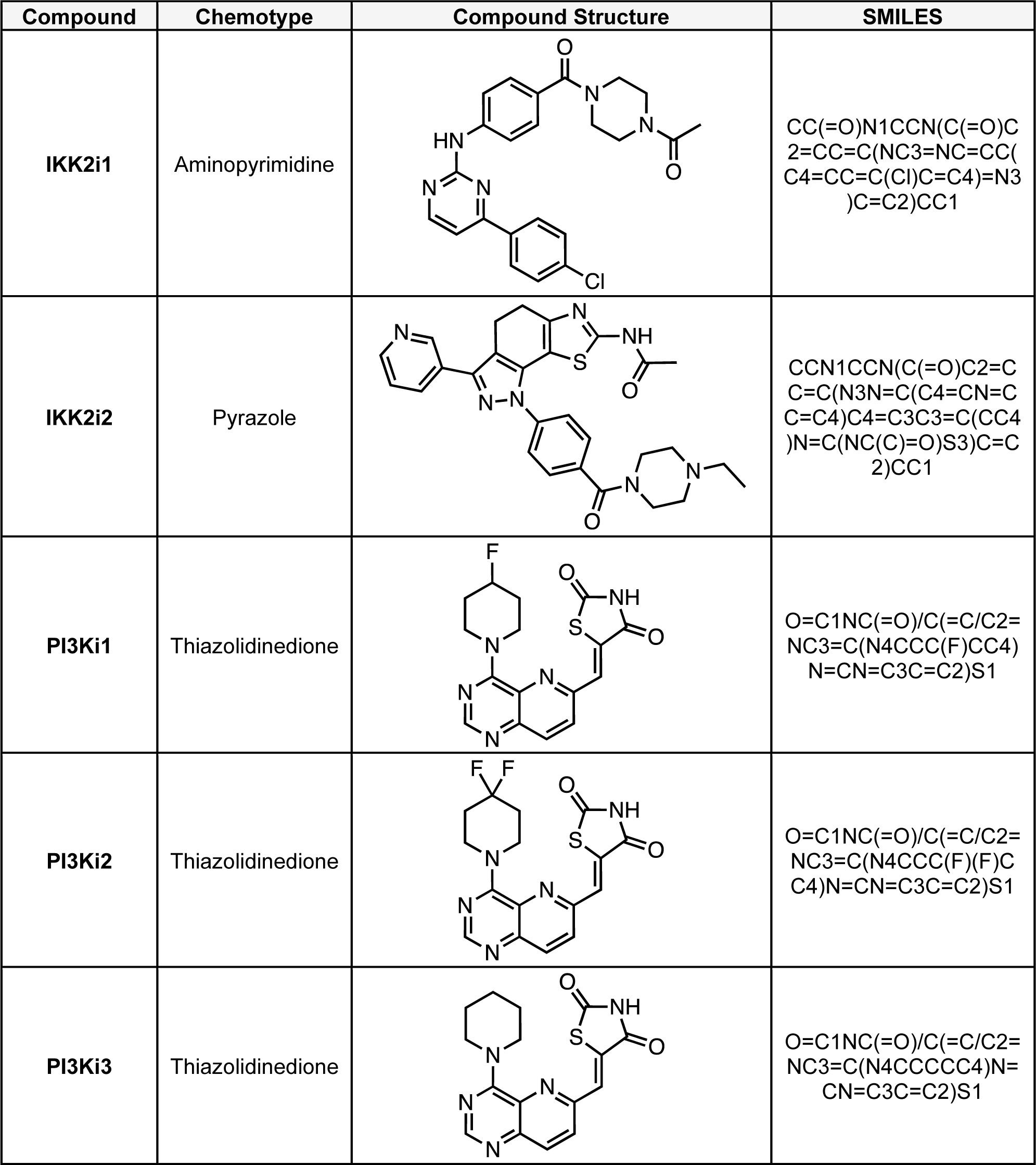
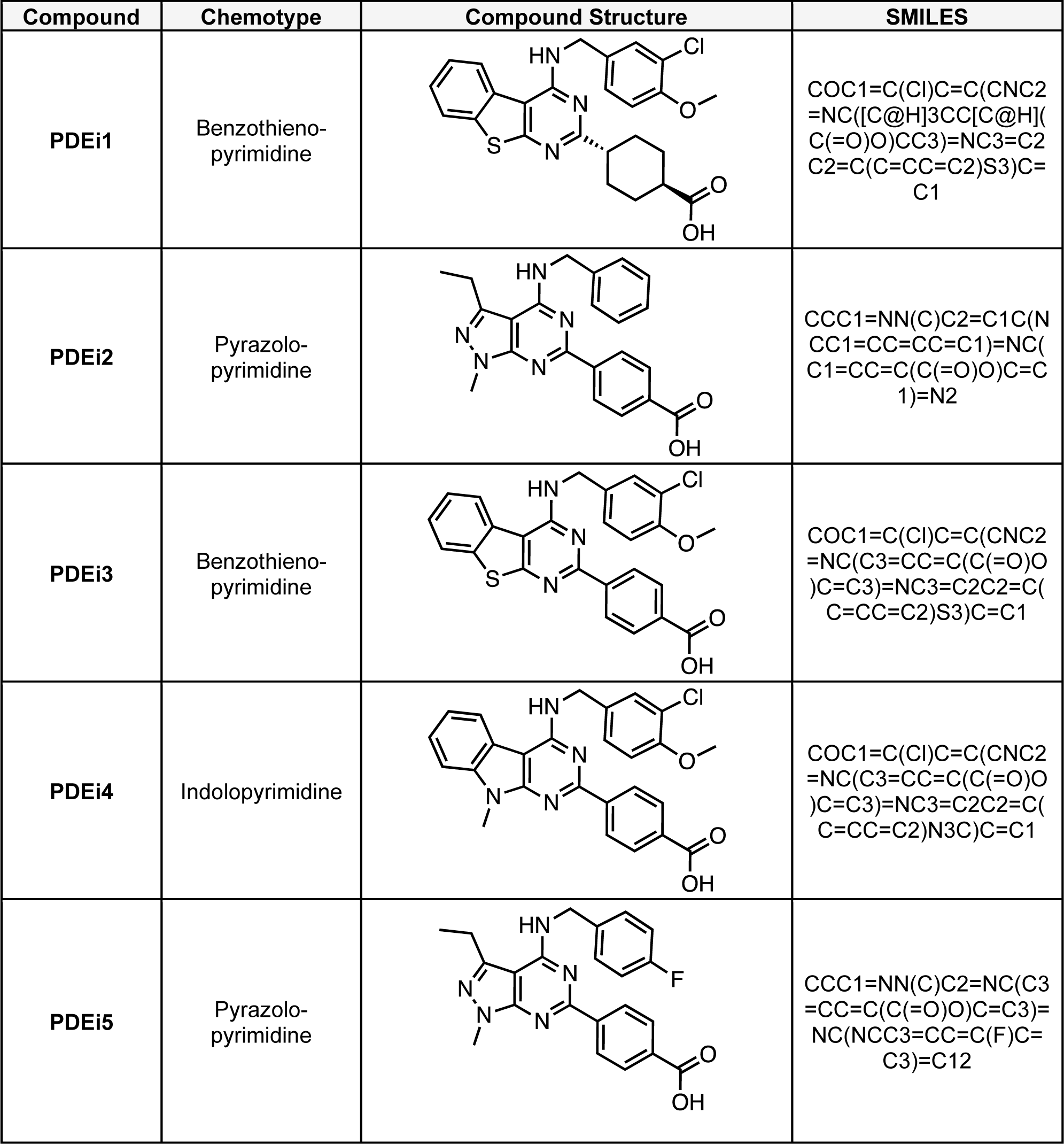
Chemical structures of compounds tested *in vivo*.

**Table S3:** *In vitro* inhibition of selected mammalian PDEs by PDEi2.

| Enzyme | Tissue | IC<sub>50</sub> (nM) |
| --- | --- | --- |
| PDE-III | Guinea pig, Heart | 1400 |
| PDE-I | Guinea pig, Heart | 600 |
| PDE-IV | Guinea pig, Heart | 1500 |
| PDE-V | Guinea pig, Lung | 20 |
| PDE-VI | Bovine, Retina | 70 |

**Table S4:**
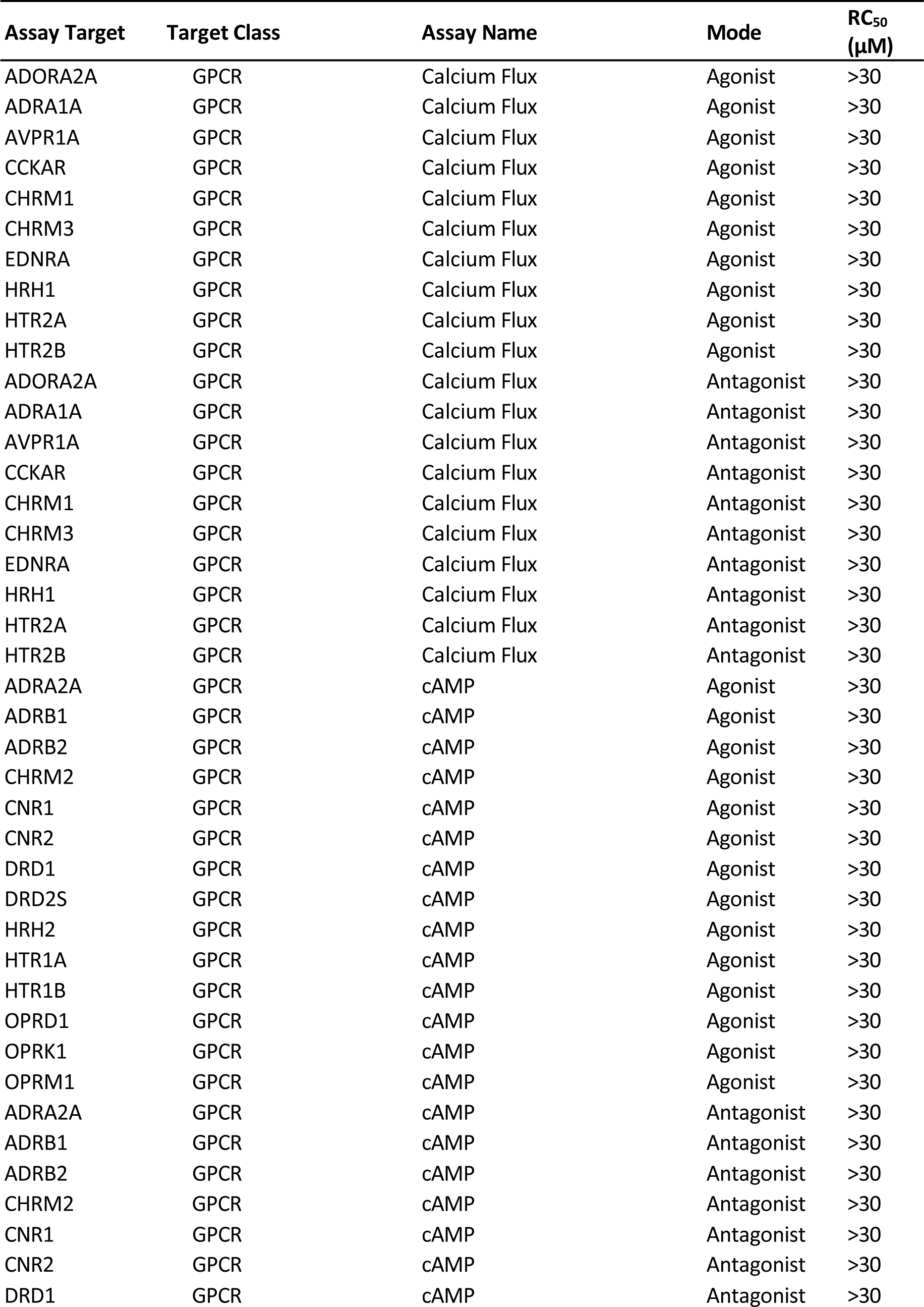
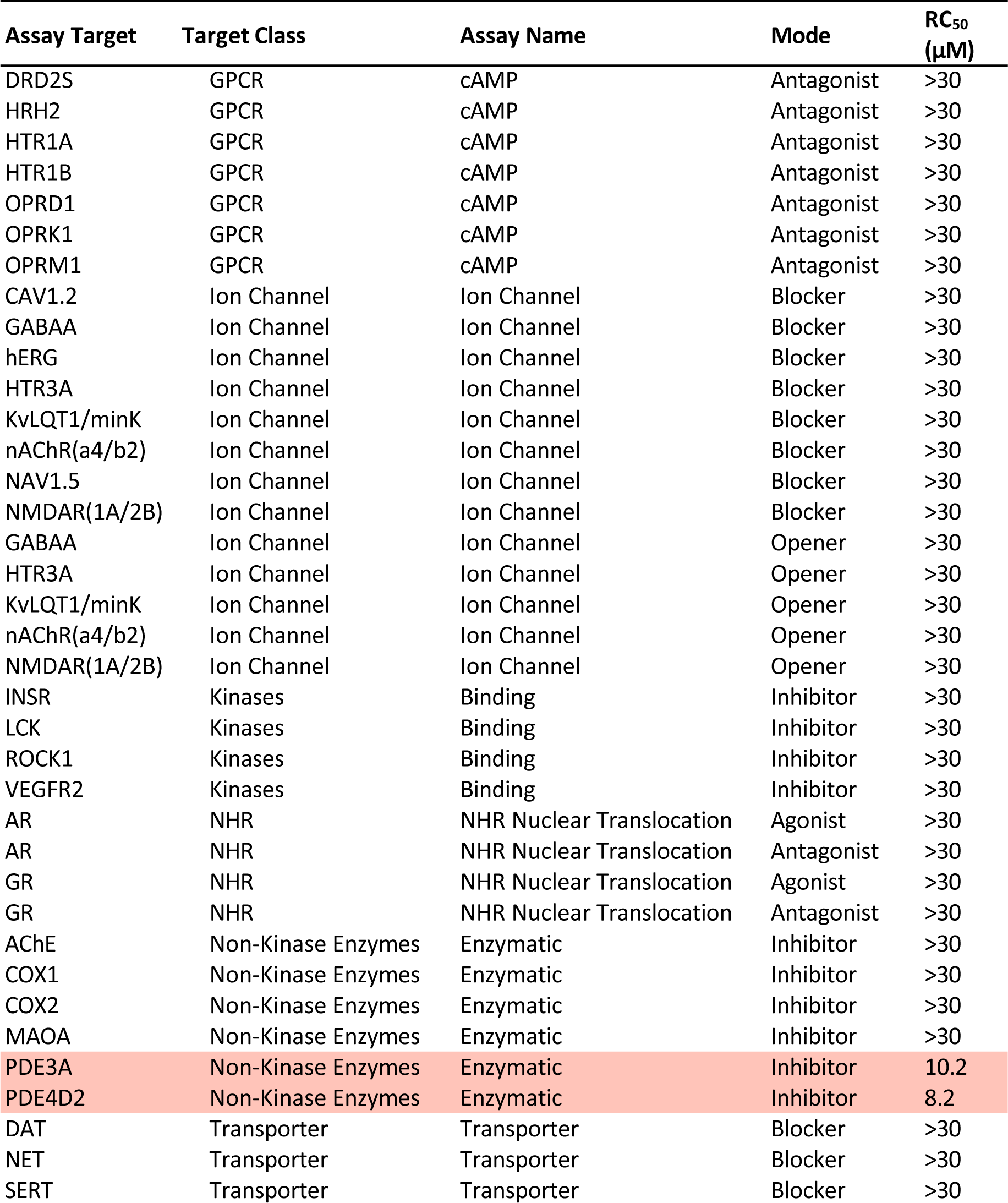
Off-targets assay results for PDEi2. Data from Eurofins SafetyScan47 assay set.

| Assay Target | Target Class | Assay Name | Mode | RC<sub>50</sub><br>(μM) |
| --- | --- | --- | --- | --- |
| ADORA2A | GPCR | Calcium Flux | Agonist | >30 |
| ADRA1A | GPCR | Calcium Flux | Agonist | >30 |
| AVPR1A | GPCR | Calcium Flux | Agonist | >30 |
| CCKAR | GPCR | Calcium Flux | Agonist | >30 |
| CHRM1 | GPCR | Calcium Flux | Agonist | >30 |
| CHRM3 | GPCR | Calcium Flux | Agonist | >30 |
| EDNRA | GPCR | Calcium Flux | Agonist | >30 |
| HRH1 | GPCR | Calcium Flux | Agonist | >30 |
| HTR2A | GPCR | Calcium Flux | Agonist | >30 |
| HTR2B | GPCR | Calcium Flux | Agonist | >30 |
| ADORA2A | GPCR | Calcium Flux | Antagonist | >30 |
| ADRA1A | GPCR | Calcium Flux | Antagonist | >30 |
| AVPR1A | GPCR | Calcium Flux | Antagonist | >30 |
| CCKAR | GPCR | Calcium Flux | Antagonist | >30 |
| CHRM1 | GPCR | Calcium Flux | Antagonist | >30 |
| CHRM3 | GPCR | Calcium Flux | Antagonist | >30 |
| EDNRA | GPCR | Calcium Flux | Antagonist | >30 |
| HRH1 | GPCR | Calcium Flux | Antagonist | >30 |
| HTR2A | GPCR | Calcium Flux | Antagonist | >30 |
| HTR2B | GPCR | Calcium Flux | Antagonist | >30 |
| ADRA2A | GPCR | cAMP | Agonist | >30 |
| ADRB1 | GPCR | cAMP | Agonist | >30 |
| ADRB2 | GPCR | cAMP | Agonist | >30 |
| CHRM2 | GPCR | cAMP | Agonist | >30 |
| CNR1 | GPCR | cAMP | Agonist | >30 |
| CNR2 | GPCR | cAMP | Agonist | >30 |
| DRD1 | GPCR | cAMP | Agonist | >30 |
| DRD2S | GPCR | cAMP | Agonist | >30 |
| HRH2 | GPCR | cAMP | Agonist | >30 |
| HTR1A | GPCR | cAMP | Agonist | >30 |
| HTR1B | GPCR | cAMP | Agonist | >30 |
| OPRD1 | GPCR | cAMP | Agonist | >30 |
| OPRK1 | GPCR | cAMP | Agonist | >30 |
| OPRM1 | GPCR | cAMP | Agonist | >30 |
| ADRA2A | GPCR | cAMP | Antagonist | >30 |
| ADRB1 | GPCR | cAMP | Antagonist | >30 |
| ADRB2 | GPCR | cAMP | Antagonist | >30 |
| CHRM2 | GPCR | cAMP | Antagonist | >30 |
| CNR1 | GPCR | cAMP | Antagonist | >30 |
| CNR2 | GPCR | cAMP | Antagonist | >30 |
| DRD1 | GPCR | cAMP | Antagonist | >30 |

| Assay Target | Target Class | Assay Name | Mode | RC<sub>50</sub><br>(μM) |
| --- | --- | --- | --- | --- |
| DRD2S | GPCR | cAMP | Antagonist | >30 |
| HRH2 | GPCR | cAMP | Antagonist | >30 |
| HTR1A | GPCR | cAMP | Antagonist | >30 |
| HTR1B | GPCR | cAMP | Antagonist | >30 |
| OPRD1 | GPCR | cAMP | Antagonist | >30 |
| OPRK1 | GPCR | cAMP | Antagonist | >30 |
| OPRM1 | GPCR | cAMP | Antagonist | >30 |
| CAV1.2 | Ion Channel | Ion Channel | Blocker | >30 |
| GABAA | Ion Channel | Ion Channel | Blocker | >30 |
| hERG | Ion Channel | Ion Channel | Blocker | >30 |
| HTR3A | Ion Channel | Ion Channel | Blocker | >30 |
| KvLQT1/minK | Ion Channel | Ion Channel | Blocker | >30 |
| nAChR(α4/β2) | Ion Channel | Ion Channel | Blocker | >30 |
| NAV1.5 | Ion Channel | Ion Channel | Blocker | >30 |
| NMDAR(1A/2B) | Ion Channel | Ion Channel | Blocker | >30 |
| GABAA | Ion Channel | Ion Channel | Opener | >30 |
| HTR3A | Ion Channel | Ion Channel | Opener | >30 |
| KvLQT1/minK | Ion Channel | Ion Channel | Opener | >30 |
| nAChR(α4/β2) | Ion Channel | Ion Channel | Opener | >30 |
| NMDAR(1A/2B) | Ion Channel | Ion Channel | Opener | >30 |
| INSR | Kinases | Binding | Inhibitor | >30 |
| LCK | Kinases | Binding | Inhibitor | >30 |
| ROCK1 | Kinases | Binding | Inhibitor | >30 |
| VEGFR2 | Kinases | Binding | Inhibitor | >30 |
| AR | NHR | NHR Nuclear Translocation | Agonist | >30 |
| AR | NHR | NHR Nuclear Translocation | Antagonist | >30 |
| GR | NHR | NHR Nuclear Translocation | Agonist | >30 |
| GR | NHR | NHR Nuclear Translocation | Antagonist | >30 |
| AChE | Non-Kinase Enzymes | Enzymatic | Inhibitor | >30 |
| COX1 | Non-Kinase Enzymes | Enzymatic | Inhibitor | >30 |
| COX2 | Non-Kinase Enzymes | Enzymatic | Inhibitor | >30 |
| MAOA | Non-Kinase Enzymes | Enzymatic | Inhibitor | >30 |
| PDE3A | Non-Kinase Enzymes | Enzymatic | Inhibitor | 10.2 |
| PDE4D2 | Non-Kinase Enzymes | Enzymatic | Inhibitor | 8.2 |
| DAT | Transporter | Transporter | Blocker | >30 |
| NET | Transporter | Transporter | Blocker | >30 |
| SERT | Transporter | Transporter | Blocker | >30 |

**Table S5:** Inhibition of human phosphodiesterases by PDEi5 at 10 µM. Assays performed by Eurofins. Enzymes inhibited >50% at 10 µM PDEi5 highlighted in red.

| Enzyme | Enzyme Source | Substrate | % Inhibition relative to control |  |  |
| --- | --- | --- | --- | --- | --- |
|  |  |  | Replicate 1 | Replicate 2 | Mean |
| **PDE1B** | human recombinant (Sf9 cells) | [3H]cGMP | 19.1 | 19.1 | 18 |
| **PDE2A1** | human recombinant (Sf21 cells) | [3H]cAMP | 35.7 | 35.7 | 36.4 |
| **PDE3A** | human recombinant (Sf21 cells) | [3H]cAMP | 30.1 | 30.1 | 27.4 |
| **PDE3B** | human recombinant (Sf9 cells) | [3H]cAMP | -0.4 | -0.4 | 2 |
| **PDE4A1A** | human recombinant (Sf9 cells) | [3H]cAMP | 27.7 | 27.7 | 31.2 |
| **PDE4B1** | human recombinant (Sf9 cells) | [3H]cAMP | 49 | 49 | 54.3 |
| **PDE4D2** | human recombinant (Sf9 cells) | [3H]cAMP | 42.1 | 42.1 | 42 |
| **PDE5 (non-selective)** | human platelets | [3H]cGMP | 91.5 | 91.5 | 92.9 |
| **PDE6 (non-selective)** | bovine retina | [3H]cGMP | 86.9 | 86.9 | 79.3 |
| **PDE7A1** | human recombinant (Sf9 cells) | [3H]cAMP | 23.2 | 23.2 | 21.6 |
| **PDE8A1** | human recombinant (Sf9 cells) | [3H]cAMP | 32.9 | 32.9 | 30 |
| **PDE10A2** | human recombinant (Sf9 cells) | [3H]cAMP | 32.2 | 32.2 | 36.5 |
| **PDE11A** | human recombinant (Sf9 cells) | [3H]cAMP | 59.1 | 59.1 | 55.3 |

**Table S6:** Inhibition of recombinant human CYPS by PDEi5 at 10 µM. Data generated by Eurofins.

| Enzyme | % Inhibition relative to control |  |  |
| --- | --- | --- | --- |
|  | Replicate 1 | Replicate 2 | Mean |
| **CYP1A2** | 3.1 | 20.3 | 11.7 |
| **CYP2B6** | 2.8 | 4.2 | 3.5 |
| **CYP2C8** | 34.7 | 28.1 | 31.4 |
| **CYP2C9** | 20.8 | 15.6 | 18.2 |
| **CYP2C19** | 0.6 | -8 | -3.7 |
| **CYP2D6** | 7.9 | 11.5 | 9.7 |
| **CYP3A4** | 6.3 | 7 | 6.6 |

## References

1 Checkley, W. et al. A review of the global burden, novel diagnostics, therapeutics, and vaccine targets for cryptosporidium. Lancet Infect Dis 15, 85–94 (2015). 10.1016/S1473-3099(14)70772-8

2 Hlavsa, M. C. et al. Surveillance for waterborne disease outbreaks and other health events associated with recreational water --- United States, 2007--2008. MMWR Surveill Summ 60, 1–32 (2011).

3 Liu, J. et al. Use of quantitative molecular diagnostic methods to identify causes of diarrhoea in children: a reanalysis of the GEMS case-control study. Lancet 388, 1291–1301 (2016). 10.1016/S0140-6736(16)31529-X

4 Khalil, I. A. et al. Morbidity, mortality, and long-term consequences associated with diarrhoea from Cryptosporidium infection in children younger than 5 years: a meta-analyses study. Lancet Glob Health 6, e758–e768 (2018). 10.1016/S2214-109X(18)30283-3

5 Amadi, B. et al. Effect of nitazoxanide on morbidity and mortality in Zambian children with cryptosporidiosis: a randomised controlled trial. Lancet 360, 1375–1380 (2002). 10.1016/S0140-6736(02)11401-2

6 Amadi, B. et al. High dose prolonged treatment with nitazoxanide is not effective for cryptosporidiosis in HIV positive Zambian children: a randomised controlled trial. BMC Infect Dis 9, 195 (2009). 10.1186/1471-2334-9-195

7 Bessoff, K. et al. Identification of Cryptosporidium parvum active chemical series by Repurposing the open access malaria box. Antimicrob Agents Chemother 58, 2731–2739 (2014). 10.1128/AAC.02641-13

8 Buckner, F. S. et al. Optimization of Methionyl tRNA-Synthetase Inhibitors for Treatment of Cryptosporidium Infection. Antimicrob Agents Chemother 63 (2019). 10.1128/AAC.02061-18

9 Love, M. S. et al. A high-throughput phenotypic screen identifies clofazimine as a potential treatment for cryptosporidiosis. PLoS Negl Trop Dis 11, e0005373 (2017). 10.1371/journal.pntd.0005373

10 Baragana, B. et al. Lysyl-tRNA synthetase as a drug target in malaria and cryptosporidiosis. Proc Natl Acad Sci U S A 116, 7015–7020 (2019). 10.1073/pnas.1814685116

11 Castellanos-Gonzalez, A. et al. A novel calcium-dependent protein kinase inhibitor as a lead compound for treating cryptosporidiosis. J Infect Dis 208, 1342–1348 (2013). 10.1093/infdis/jit327

12 Murphy, R. C. et al. Discovery of Potent and Selective Inhibitors of Calcium-Dependent Protein Kinase 1 (CDPK1) from C. parvum and T. gondii. ACS Med Chem Lett 1, 331–335 (2010). 10.1021/ml100096t

13 Schaefer, D. A. et al. Novel Bumped Kinase Inhibitors Are Safe and Effective Therapeutics in the Calf Clinical Model for Cryptosporidiosis. J Infect Dis 214, 1856–1864 (2016). 10.1093/infdis/jiw488

14 Guo, F. et al. Amelioration of Cryptosporidium parvum infection in vitro and in vivo by targeting parasite fatty acyl-coenzyme A synthetases. J Infect Dis 209, 1279–1287 (2014). 10.1093/infdis/jit645

15 Guo, F., Zhang, H., McNair, N. N., Mead, J. R. & Zhu, G. The Existing Drug Vorinostat as a New Lead Against Cryptosporidiosis by Targeting the Parasite Histone Deacetylases. J Infect Dis 217, 1110–1117 (2018). 10.1093/infdis/jix689

16 Jain, V. et al. Targeting Prolyl-tRNA Synthetase to Accelerate Drug Discovery against Malaria, Leishmaniasis, Toxoplasmosis, Cryptosporidiosis, and Coccidiosis. Structure 25, 1495–1505 e1496 (2017). 10.1016/j.str.2017.07.015

17 Huang, W. et al. 5-Aminopyrazole-4-Carboxamide-Based Compounds Prevent the Growth of Cryptosporidium parvum. Antimicrob Agents Chemother 61 (2017). 10.1128/AAC.00020-17

18 Manjunatha, U. H. et al. A Cryptosporidium PI(4)K inhibitor is a drug candidate for cryptosporidiosis. Nature 546, 376–380 (2017). 10.1038/nature22337

19 Lunde, C. S. et al. Identification of a potent benzoxaborole drug candidate for treating cryptosporidiosis. Nat Commun 10, 2816 (2019). 10.1038/s41467-019-10687-y

20 Jumani, R. S. et al. A Novel Piperazine-Based Drug Lead for Cryptosporidiosis from the Medicines for Malaria Venture Open-Access Malaria Box. Antimicrob Agents Chemother 62, e01505–01517 (2018). 10.1128/AAC.01505-17

21 Vinayak, S. et al. Bicyclic azetidines kill the diarrheal pathogen Cryptosporidium in mice by inhibiting parasite phenylalanyl-tRNA synthetase. Sci Transl Med 12 (2020). 10.1126/scitranslmed.aba8412

22 Oboh, E. et al. Optimization of the Urea Linker of Triazolopyridazine MMV665917 Results in a New Anticryptosporidial Lead with Improved Potency and Predicted hERG Safety Margin. J Med Chem 64, 11729–11745 (2021). 10.1021/acs.jmedchem.1c01136

23 Hasan, M. M. et al. Spontaneous Selection of Cryptosporidium Drug Resistance in a Calf Model of Infection. Antimicrob Agents Chemother 65 (2021). 10.1128/AAC.00023-21

24 Heemskerk, J., Tobin, A. J. & Bain, L. J. Teaching old drugs new tricks. Meeting of the Neurodegeneration Drug Screening Consortium, 7-8 April 2002, Washington, DC, USA. Trends Neurosci 25, 494–496 (2002). 10.1016/s0166-2236(02)02236-1

25 De Pascale, G. et al. Remdesivir plus Dexamethasone in COVID-19: A cohort study of severe patients requiring high flow oxygen therapy or non-invasive ventilation. PLoS One 17, e0267038 (2022). 10.1371/journal.pone.0267038

26 Edwards, A. What Are the Odds of Finding a COVID-19 Drug from a Lab Repurposing Screen? J Chem Inf Model 60, 5727–5729 (2020). 10.1021/acs.jcim.0c00861

27 Bessoff, K., Sateriale, A., Lee, K. K. & Huston, C. D. Drug repurposing screen reveals FDA- approved inhibitors of human HMG-CoA reductase and isoprenoid synthesis that block Cryptosporidium parvum growth. Antimicrob Agents Chemother 57, 1804–1814 (2013). 10.1128/AAC.02460-12

28 Arnold, S. L. M. et al. Necessity of Bumped Kinase Inhibitor Gastrointestinal Exposure in Treating Cryptosporidium Infection. J Infect Dis 216, 55–63 (2017). 10.1093/infdis/jix247

29 Iroh Tam, P., et al. Clofazimine for Treatment of Cryptosporidiosis in Human Immunodeficiency Virus Infected Adults: An Experimental Medicine, Randomized, Double-blind, Placebo-controlled Phase 2a Trial. Clin Infect Dis 73, 183–191 (2021). 10.1093/cid/ciaa421

30 Wermuth, C. G. Selective optimization of side activities: the SOSA approach. Drug Discov Today 11, 160–164 (2006). 10.1016/S1359-6446(05)03686-X

31 Merck-KGaA. Biopharma Mini Library, <https://www.emdgroup.com/en/research/open-innovation/biopharma-open-innovation-portal/biopharma-mini-library.html> (2023).

32. 32 Merck-KGaA. Open Global Health Library, <https://www.emdgroup.com/en/research/open-innovation/biopharma-open-innovation-portal/open-global-health-library.html> (2023).

33 Baell, J. B. & Holloway, G. A. New substructure filters for removal of pan assay interference compounds (PAINS) from screening libraries and for their exclusion in bioassays. J Med Chem 53, 2719–2740 (2010). 10.1021/jm901137j

34 Camps, M. et al. Blockade of PI3Kgamma suppresses joint inflammation and damage in mouse models of rheumatoid arthritis. Nat Med 11, 936–943 (2005). 10.1038/nm1284

35 Pomel, V. et al. Furan-2-ylmethylene thiazolidinediones as novel, potent, and selective inhibitors of phosphoinositide 3-kinase gamma. J Med Chem 49, 3857–3871 (2006). 10.1021/jm0601598

36 Maggiora, G., Vogt, M., Stumpfe, D. & Bajorath, J. Molecular similarity in medicinal chemistry. J Med Chem 57, 3186–3204 (2014). 10.1021/jm401411z

37 Burch, H. A. Nitrofuryl heterocycles. VII. 4-Amino-6-(5-nitro-2-furyl)-1H-pyrazolo[3,4- d]pyrimidines. J Med Chem 11, 79–83 (1968). 10.1021/jm00307a017

38 Mouton, J. W. et al. Tissue concentrations: do we ever learn? J Antimicrob Chemother 61, 235–237 (2008). 10.1093/jac/dkm476

39 Rawson, D. J. et al. The discovery of UK-369003, a novel PDE5 inhibitor with the potential for oral bioavailability and dose-proportional pharmacokinetics. Bioorg Med Chem 20, 498–509 (2012). 10.1016/j.bmc.2011.10.022

40 Fiorito, J. et al. Synthesis of quinoline derivatives: discovery of a potent and selective phosphodiesterase 5 inhibitor for the treatment of Alzheimer’s disease. Eur J Med Chem 60, 285–294 (2013). 10.1016/j.ejmech.2012.12.009

41 Jumper, J. et al. Highly accurate protein structure prediction with AlphaFold. Nature 596, 583–589 (2021). 10.1038/s41586-021-03819-2

42 Varadi, M. et al. AlphaFold Protein Structure Database: massively expanding the structural coverage of protein-sequence space with high-accuracy models. Nucleic Acids Res 50, D439–D444 (2022). 10.1093/nar/gkab1061

43 Tandel, J. et al. Life cycle progression and sexual development of the apicomplexan parasite Cryptosporidium parvum. Nat Microbiol 4, 2226–2236 (2019). 10.1038/s41564-019-0539-x

44 Brown, D. G. et al. (Protein Data Bank, 2018).

45 Friesner, R. A. et al. Glide: a new approach for rapid, accurate docking and scoring. 1. Method and assessment of docking accuracy. J Med Chem 47, 1739–1749 (2004). 10.1021/jm0306430

46 Jumani, R. S. et al. A suite of phenotypic assays to ensure pipeline diversity when prioritizing drug-like Cryptosporidium growth inhibitors. Nat Commun 10, 1862 (2019). 10.1038/s41467-019-09880-w

47. 47 Eurofins. *In* Vitro Safety Pharmacology Profiling Panels, <https://www.eurofinsdiscovery.com/solution/safety-panels> (2023).

48 Banerjee, P., Eckert, A. O., Schrey, A. K. & Preissner, R. ProTox-II: a webserver for the prediction of toxicity of chemicals. Nucleic Acids Res 46, W257–W263 (2018). 10.1093/nar/gky318

49 Campbell, L. D., Stewart, J. N. & Mead, J. R. Susceptibility to Cryptosporidium parvum infections in cytokine- and chemokine-receptor knockout mice. J Parasitol 88, 1014–1016 (2002). 10.1645/0022-3395(2002)088[1014:STCPII]2.0.CO;2

50 Smith, L. M., Bonafonte, M. T., Campbell, L. D. & Mead, J. R. Exogenous interleukin-12 (IL-12) exacerbates Cryptosporidium parvum infection in gamma interferon knockout mice. Exp Parasitol 98, 123–133 (2001). 10.1006/expr.2001.4627

51 Gorla, S. K. et al. Validation of IMP dehydrogenase inhibitors in a mouse model of cryptosporidiosis. Antimicrob Agents Chemother 58, 1603–1614 (2014). 10.1128/AAC.02075-13

52 Veerman, J. et al. Synthesis and evaluation of analogs of the phenylpyridazinone NPD-001 as potent trypanosomal TbrPDEB1 phosphodiesterase inhibitors and in vitro trypanocidals. Bioorg Med Chem 24, 1573–1581 (2016). 10.1016/j.bmc.2016.02.032

53 Blaazer, A. R. et al. Targeting a Subpocket in Trypanosoma brucei Phosphodiesterase B1 (TbrPDEB1) Enables the Structure-Based Discovery of Selective Inhibitors with Trypanocidal Activity. J Med Chem 61, 3870–3888 (2018). 10.1021/acs.jmedchem.7b01670

54 de Heuvel, E. et al. Alkynamide phthalazinones as a new class of TbrPDEB1 inhibitors. Bioorg Med Chem 27, 3998–4012 (2019). 10.1016/j.bmc.2019.06.027

55 Amata, E., Bland, N. D., Campbell, R. K. & Pollastri, M. P. Evaluation of pyrrolidine and pyrazolone derivatives as inhibitors of trypanosomal phosphodiesterase B1 (TbrPDEB1). Tetrahedron Lett 56, 2832–2835 (2015). 10.1016/j.tetlet.2015.04.061

56 Zheng, Y. et al. Discovery of 5-Phenylpyrazolopyrimidinone Analogs as Potent Antitrypanosomal Agents with In Vivo Efficacy. J Med Chem 66, 10252–10264 (2023). 10.1021/acs.jmedchem.3c00161

57 Long, T. et al. Phenotypic, chemical and functional characterization of cyclic nucleotide phosphodiesterase 4 (PDE4) as a potential anthelmintic drug target. PLoS Negl Trop Dis 11, e0005680 (2017). 10.1371/journal.pntd.0005680

58 Sebastian-Perez, V. et al. Discovery of novel Schistosoma mansoni PDE4A inhibitors as potential agents against schistosomiasis. Future Med Chem 11, 1703–1720 (2019). 10.4155/fmc-2018-0592

59 Zheng, Y. et al. To Target or Not to Target Schistosoma mansoni Cyclic Nucleotide Phosphodiesterase 4A? Int J Mol Sci 24 (2023). 10.3390/ijms24076817

60 Botros, S. S. et al. The phosphodiesterase-4 inhibitor roflumilast impacts Schistosoma mansoni ovipositing in vitro but displays only modest antischistosomal activity in vivo. Exp Parasitol 208, 107793 (2020). 10.1016/j.exppara.2019.107793

61 Howard, B. L. et al. Identification of potent phosphodiesterase inhibitors that demonstrate cyclic nucleotide-dependent functions in apicomplexan parasites. ACS Chem Biol 10, 1145–1154 (2015). 10.1021/cb501004q

62 Zheng, Y. et al. Structural Optimization of BIPPO Analogs as Potent Antimalarials. Molecules 28 (2023). 10.3390/molecules28134939

63 Gut, J. & Nelson, R. G. Cryptosporidium parvum: synchronized excystation in vitro and evaluation of sporozoite infectivity with a new lectin-based assay. J Eukaryot Microbiol 46, 56S–57S (1999).

64 Parr, J. B. et al. Detection and quantification of Cryptosporidium in HCT-8 cells and human fecal specimens using real-time polymerase chain reaction. Am J Trop Med Hyg 76, 938–942 (2007).

65 Amos, B. et al. VEuPathDB: the eukaryotic pathogen, vector and host bioinformatics resource center. Nucleic Acids Res 50, D898–D911 (2022). 10.1093/nar/gkab929

66 Altschul, S. F., Gish, W., Miller, W., Myers, E. W. & Lipman, D. J. Basic local alignment search tool. J Mol Biol 215, 403–410 (1990). 10.1016/S0022-2836(05)80360-2

